# Anteroposterior patterning of the zebrafish ear through Fgf- and Hh-dependent regulation of *hmx3a* expression

**DOI:** 10.1101/451963

**Authors:** Ryan D. Hartwell, Samantha J. England, Nicholas A. M. Monk, Nicholas J. van Hateren, Sarah Baxendale, Mar Marzo, Katharine E. Lewis, Tanya T. Whitfield

## Abstract

In the zebrafish, Fgf and Hh signalling assign anterior and posterior identity, respectively, to the poles of the developing ear. Mis-expression of *fgf3* or inhibition of Hh signalling results in double-anterior ears, including ectopic expression of *hmx3a*. To understand how this double-anterior pattern is established, we characterised transcriptional responses in Fgf gain-of-signalling or Hh loss-of-signalling backgrounds. Mis-expression of *fgf3* resulted in rapid expansion of anterior otic markers, refining over time to give the duplicated pattern. Response to Hh inhibition was very different: initial anteroposterior asymmetry was retained, with de novo duplicate expression domains appearing later. We show that Hmx3a is required for normal anterior otic patterning, but neither loss nor gain of *hmx3a* function was sufficient to generate ear duplications. Using our data to infer a transcriptional regulatory network required for acquisition of otic anterior identity, we can recapitulate both the wild-type and the double-anterior pattern in a mathematical model.

## Introduction

The otic placode—precursor of the vertebrate inner ear—has the remarkable ability to generate a mirror-image organ with duplicate structures under some experimental conditions in fish and amphibians, as originally described by R. G. Harrison over eighty years ago (reviewed in (Whitfield and Hammond, 2007)). Understanding the generation of such duplicated structures can give us fundamental insights into mechanisms of organ patterning, tissue polarity and symmetry-breaking during embryogenesis. During normal development in the zebrafish, anteroposterior asymmetries in otic gene expression are evident as early as the 4-somite stage (11.5 hours post fertilisation (hpf)), when expression of the transcription factor gene *hmx3a* appears at the anterior of the otic placode (Feng and Xu, 2010). Additional genes with predominantly anterior patterns of expression in the otic placode or vesicle begin to be expressed over the next 10 hours, including the transcription factor genes *hmx2* and *pax5* (Feng and Xu, 2010; Kwak et al., 2006), together with the fibroblast growth factor (Fgf) family genes *fgf3, fgf8a* and *fgf10a* (Léger and Brand, 2002; McCarroll and Nechiporuk, 2013). Later, at otic vesicle stages (24 hpf onwards), the size and position of the otoliths, together with the position, shape and planar polarity patterns of the sensory maculae, provide landmarks for distinguishing anterior and posterior structures in the ear (Hammond and Whitfield, 2011) (Fig. 1). In addition, a few markers begin to be expressed specifically in posterior otic tissue (*pou3f3b, bmp7a* and *fsta*) at otic vesicle stages (Kwak et al., 2006; Mowbray et al., 2001; Schmid et al., 2000), but these are not reliable posterior markers at earlier otic placode stages.

**Figure 1.**
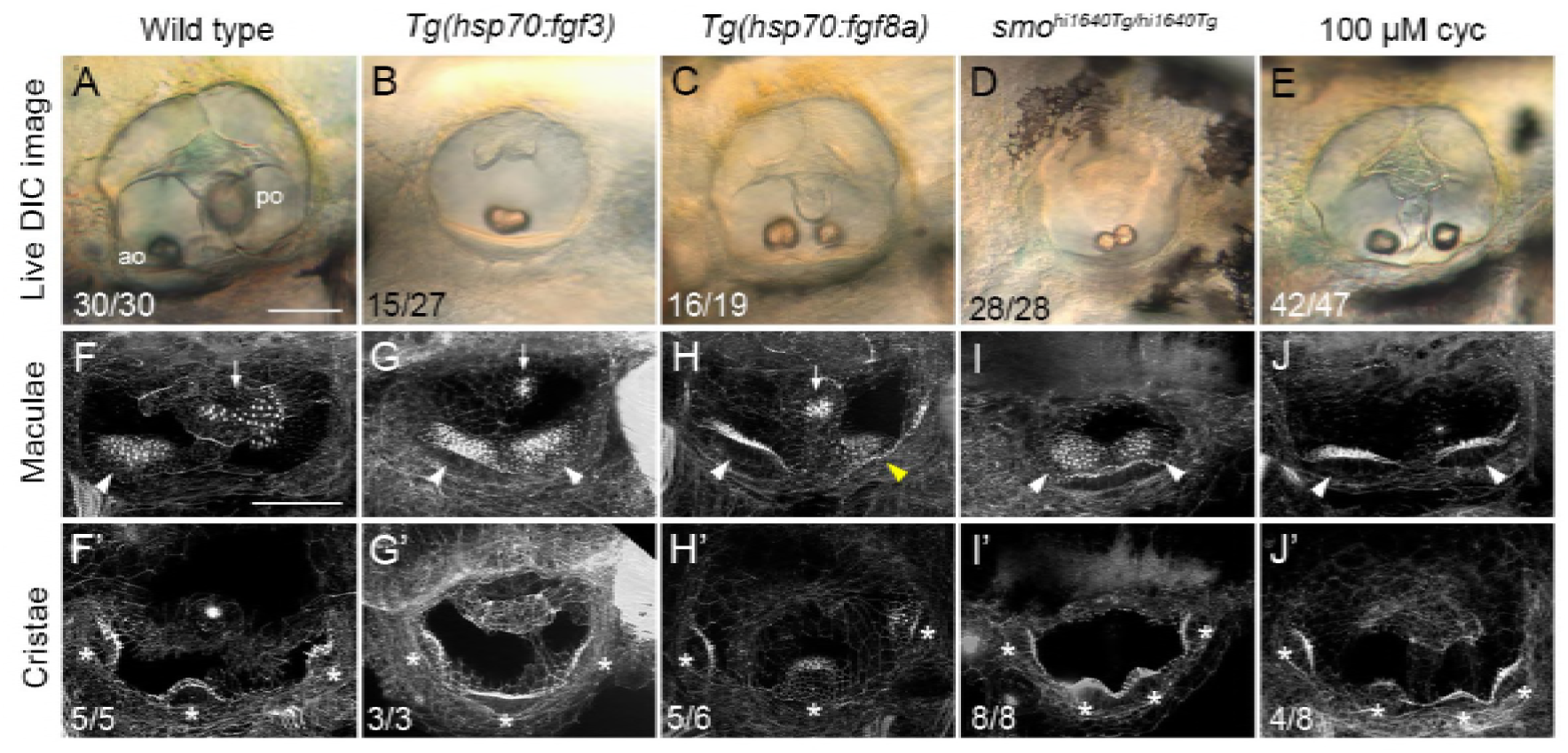
Duplicate double-anterior ear phenotypes resulting from early Fgf mis-expression or Hh pathway inhibition. **(A-E)** Differential interference contrast (DIC) images of ears in live embryos at 3 dpf (72 hpf). **(F-J’)** Confocal images of FITC-phalloidin stains, revealing stereociliary bundles on sensory hair cells in the maculae (F-J) or cristae (F’—J’). Anterior maculae and duplicate anterior maculae are marked with arrowheads; posterior maculae and remnants of posterior maculae are marked with arrows. Cristae and duplicate cristae are marked with asterisks. Yellow arrowhead in H indicates macula that is ventromedial in position, and close to remnants of the posterior macula (arrowhead). Note the enlarged lateral crista in G’. (The bright spot in the centre of F’ is a lateral line neuromast.) Representative phenotypes are shown; numbers of embryos displaying these phenotypes are indicated on the panels. All *Tg(hsp70:fgf3)* heat-shocked embryos (*n*=27) and *smo^hl1640Tg/hl1640Tg^* mutants (*n*=28) showed double-anterior ears. In B, 15/27 ears had a single fused otolith as shown; the remaining 12/27 ears had two separate, but small and ventrally-positioned otoliths. In D, 17/28 ears had two otoliths touching as shown; in the remaining 11/28 ears, the otoliths were separate, but both small and ventrally positioned. Genotypes or treatments are indicated for each column. Transgenic lines were subject to 30 minutes of heat shock at 14 hpf. Additional controls for this figure are shown in Figure 1—Supplemental file 1. Lateral views; anterior to the left. Abbreviations: ao, anterior (utricular) otolith; po, posterior (saccular) otolith; cyc, cyclopamine. Scale bar in A, 50 μm (applies to A—E); scale bar in F, 50 μm (applies to F—J’).

Concomitant with the appearance of anteroposterior asymmetry in the zebrafish otic domain, other early patterning events occur that are symmetrical about the anteroposterior axis. Of relevance for our study, a single sensory-competent domain, marked by the expression of *atoh1b*, splits into two domains, one at each pole of the ear, by 12 hpf. This process is dependent on Notch signalling and *atoh1b* function, and defines differences between the poles of the otic placode and a central zone (Millimaki et al., 2007). The two poles express various markers symmetrically, including *atoh1a* and *deltaD*, between 14–18 hpf (Millimaki et al., 2007), presaging the appearance of pairs of *myo7aa*-positive sensory hair cells (tether cells) at each pole by 18–24 hpf (Ernest et al., 2000). Thus, by the completion of otic induction at 14 hpf (10 somites), the otic domain has two clear poles defined by the symmetric expression of *atoh1a* and *deltaD*, with the anterior pole distinguished from the posterior by the asymmetric expression of *hmx3a*.

Although anteroposterior asymmetries in otic gene expression are already apparent by 12 hpf in the zebrafish, these can be disrupted by interfering with extrinsic signalling pathway activity after this time. For example, manipulations of either Fibroblast growth factor (Fgf) or Hedgehog (Hh) signalling between 14-19 hpf can result in striking double-anterior or double-posterior mirror-image ears. Fgf signalling is both required and sufficient to act as an anteriorising cue, whereas Hh signalling is both required and sufficient for the acquisition of posterior otic identity (Hammond et al., 2003; Hammond et al., 2010; Hammond and Whitfield, 2011). In these studies, we showed that transient *fgf3* mis-expression at 14 hpf or Hh pathway loss-of-function result in the loss of posterior-specific expression domains of *fsta* at 30 hpf and *otx1b* at 45–48 hpf, and the gain of anterior-specific gene expression in the posterior of the ear (*hmx2* and *pax5* at 24 hpf after *fgf3* mis-expression; *hmx3a* at 30 hpf in Hh loss-of-function mutants). These findings suggest that Fgf and Hh signalling normally act to establish and determine the asymmetric expression of marker genes within the otic epithelium. However, the details of their temporal mode of action in the duplication of anterior otic fates have not been explored.

In this study, we have compared the dynamics of the transcriptional responses that precede the acquisition of a duplicated anterior otic fate in an Fgf gain-of-signalling or a Hh loss-of-signalling context. Although the final duplicated ear structures appear similar after each manipulation, the early transcriptional responses differ for each signalling pathway, progressing in distinct ways to give rise to the double-anterior pattern at larval stages. One gene that shows an early transcriptional response in the zebrafish otic placode to disruption of either Fgf or Hh signalling is the *Hmx* family homeobox gene *hmx3a*. We have examined the effects of both loss-of-function and gain-of-function of *hmx3a* on inner ear patterning. Our data suggest that *hmx3a* is a key early target for the otic anteriorising activity of Fgf signalling, and that the function of *hmx3a* is required for the anterior-specific otic expression of *fgf3* and *pax5*, together with correct positioning and development of the sensory maculae. However, unlike high Fgf levels or low Hh pathway activity, mis-expression of *hmx3a* was unable to generate full duplications of anterior character at the posterior of the ear. A mathematical model based on our experimental findings can recapitulate both the wild-type and duplicated anterior pattern, allowing us to explore the dynamical principles underlying the generation of a mirror-image duplicated organ system.

## Results

### Early mis-expression of *fgf3*, but not *fgf8a*, can generate complete double-anterior ear duplications

To establish optimal conditions for generating a double-anterior ear in the zebrafish embryo, we compared otic phenotypes in transgenic lines for two different *fgf* genes, *fgf3* and *fgf8a*, with systemic transgene expression driven under the control of the *hsp70* heat-shock promoter (Lecaudey et al., 2008; Millimaki et al., 2010). Previously, we showed that a 2-hour heat shock in the *Tg(hsp70:fgf3)* line at 14 hpf (10-somite stage) resulted in a robust duplication of anterior otic structures (Hammond and Whitfield, 2011). We chose this time point to avoid any interference with otic placode induction, which is also Fgf-dependent, but is complete by 14 hpf (Kimmel et al., 1995; Phillips et al., 2001). The 14 hpf time point is also after completion of the Notch-dependent signalling event that distinguishes the otic poles from a central zone of epithelium (Millimaki et al., 2007). For the treatments described here, we reduced the time of heat shock to 30 minutes at 39°C. This shorter heat shock still results in a full ear duplication, but should minimise effects of Fgf mis-expression on other developing organ systems. After heat shock, embryos were then cultured at 33°C for a further 30 minutes, to reduce the incubation temperature gradually, before being returned to 28.5°C and incubated until 3 days post fertilisation (dpf) for processing and analysis (Fig. 1). This stepwise reduction in temperature is thought to extend transgene activation and reduce cell death following heat shock (Padanad et al., 2012; Zou et al., 1998). Non-transgenic sibling embryos, subjected to the same heat-shock treatment, served as controls (Fig. 1—Supplemental file 1).

In *Tg(hsp70:fgf3)* embryos, a 30-minute heat shock at 14 hpf gave a robust and complete duplication of anterior otic patterning at 72 hpf, as indicated by the mis-positioning or fusion of the posterior otolith, loss of posterior elements of the saccular macula, and a duplication of anterior (utricular)-like sensory elements on the posteroventral floor of the ear (Fig. 1B, G,G’). The phenotypes seen after mis-expression of *fgf8a* (30-minute heat shock at 14 hpf) were milder and more variable than those for *fgf3*, and included a split and mis-positioned saccular macula rather than a complete duplication of anterior elements (Fig. 1C, H), and a normal complement of three cristae (Fig. 1H’). A 30-minute heat shock of either transgenic line at a later stage (18 hpf) resulted in only mild effects on ear size and shape, and otolith position (Fig. 1—Supplemental file 2A–D) (Hammond and Whitfield, 2011). We therefore chose to use the *Tg(hsp70:fgf3)* line, with a 30-minute heat shock (39°C) at 14 hpf followed by 30 minutes at 33°C, in subsequent heat-shock experiments.

### Genetic or pharmacological inhibition of Hh signalling can also result in complete double-anterior ear duplications

To optimise our protocols for generating double-anterior duplicated ears through inhibition of Hh signalling, we first examined the ear phenotype in *smo*^*hi1640Tg/hi1640Tg*^ mutants. The *hi1640Tg* allele (a transgenic insertion in the *smoothened* gene, and a likely null (Chen et al., 2001)) is thought to result in a stronger reduction in Hh signalling than the point mutation alleles *smo^b641^* and *smo^b577^*, both of which predict single amino acid substitutions (Varga et al., 2001), and which we used in previous studies (Hammond et al., 2003; Hammond and Whitfield, 2011). The *smo^hi1640Tg/hi1640Tg^* mutants showed a fully penetrant double-anterior duplicated ear phenotype, with two similar-sized small otoliths located ventrally, complete loss of the posterior (saccular) macula, and duplication of the anterior (utricular) macula at the posterior of the ear, with anterior and posterior elements sometimes present as a contiguous patch of hair cells covering the ventral floor (Fig. 1D,I). Four cristae, rather than the usual three, were present in all (8/8) mutant ears imaged (Fig. 1I’). (For comparison, four cristae were present in only about 50% of ears of *smo^b641/b641^* mutant embryos (Hammond et al., 2003).)

Pharmacological inhibition of the transducer of the Hh pathway, Smoothened, using the small molecule cyclopamine, can also produce double-anterior ear duplications (Hammond et al., 2010; Sapède and Pujades, 2010). This approach enables a conditional inhibition of Hh signalling over a defined time window. For the experiments described here, we treated wild-type embryos with 100 µM cyclopamine from 14–22.5 hpf. To examine later stages, we washed out the cyclopamine at 22.5 hpf and allowed embryos to develop further until 3 dpf (72 hpf), when they were fixed for staining and imaging. Stage-matched sibling embryos—either untreated, or treated with vehicle (ethanol) only—served as controls. This cyclopamine treatment regime was sufficient to generate the double-anterior ear phenotype, characterised by two ventrally-positioned, small (utricular-like) otoliths, loss of the posterior (saccular) macula, and a clear duplication of the anterior (utricular) macula (Fig. 1E,J). Ears in 4/8 treated embryos had four cristae (Fig. 1J’); the remaining 4/8 had the normal complement of three cristae. The size and shape of the ear were less affected than in the *smo^hi1640Tg/hi1640Tg^* mutant embryos, presumably due to the transient nature of the cyclopamine treatment. Taken together, these data show that either genetic or pharmacological inhibition of Hh signalling in wild-type zebrafish embryos between 14–22.5 hpf results in a robust and reproducible double-anterior ear phenotype at 3 dpf.

### Following early *fgf3* mis-expression, otic expression of anterior markers is initially broad, with *pax5* resolving into two discrete domains

One of the most striking transcriptional changes in response to *fgf3* mis-expression is the expansion or duplication of the expression of the anterior otic markers *hmx2* and *pax5* by 24 hpf (Hammond and Whitfield, 2011). To examine the temporal dynamics of this transcriptional response, we assayed for expression of these and additional anterior otic marker genes following our optimised ‘early’ heat-shock regime (14 hpf, 30 min, 39°C) at three different time points: 16 hpf (2 hours post heat shock, to examine any rapid response), 22.5 hpf (8.5 hours post heat shock, when anterior otic expression of *hmx2* and *pax5* is strongly established in wild-type embryos) and 36 hpf (22 hours post heat shock, to examine whether any disruption to the expression pattern resolves or changes over time). For *hmx2* and *pax5*, which showed dynamic expression changes, we subsequently included two additional time points (25.5 hpf and 30 hpf) to capture these changes in more detail.

We first tested expression of three genes coding for transcription factors (*hmx3a, hmx2* and *pax5*; Fig. 2). At the earliest time point (16 hpf, two hours after heat shock), *hmx3a* showed the strongest response: expression had already expanded to cover the entire anteroposterior extent of the otic placode (Fig. 2A,B). The anterior markers *hmx2* and *pax5*, not normally expressed at this stage in wild-type embryos, were expressed at very low levels in the anterior of the otic placode of heat-shocked embryos (Fig. 2C–F). We were also able to detect widespread and robust up-regulation of the Fgf target gene *etv4* (formerly *pea3*) in transgenic embryos at this time point (Fig. 2—Supplemental file 1). By 22.5 hpf, all three transcription factor genes were strongly expressed in a broad zone across the entire anteroposterior axis of the otic vesicle in heat-shocked embryos, on the medial side, as can be seen in a dorsal view (Fig. 2G–L). Note that the overall size and shape of these otic vesicles were relatively normal. Although there was some variability (including between both ears of the same fish), the vesicles were oval in shape, indicating that otic induction had not been compromised (compare with the small, rounded vesicles of *fgf8a^ti282/ti282^* mutants, in which otic induction is disrupted (Léger and Brand, 2002)). By 36 hpf, wild-type otic expression of *hmx* genes was more complex, but a clear difference between anterior expressing and posterior non-expressing regions was evident in ventral otic epithelium (Fig. 2M, M’,O,O’). By contrast, in heat-shocked embryos, expression of *hmx3a* remained strong across the entire anteroposterior axis of the ear in ventral regions (Fig. 2N, N’); expression of *hmx2* weakened in central regions during intermediate stages, but at 36 hpf was present in a contiguous ventral domain (Fig. 2P, P’, Fig. 2—Supplemental File 2), while expression of *pax5* was lost in central regions, resolving into two discrete ventral domains at the anterior and posterior poles by 25.5 hpf (Fig. 2R, R’, Fig. 2—Supplemental File 3).

**Figure 2.**
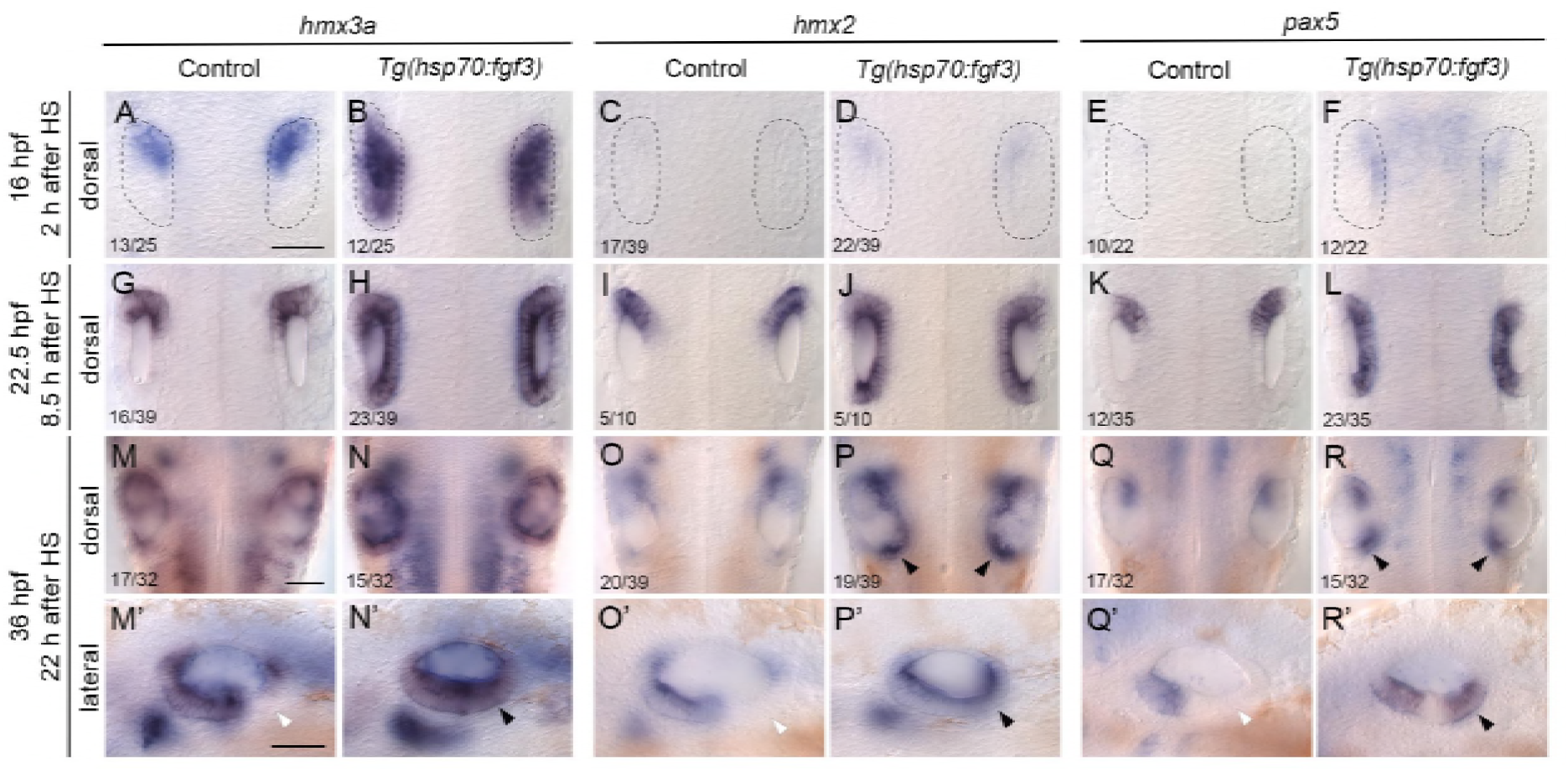
Expression of the otic anterior marker genes *hmx3a, hmx2* and *pax5* after early *fgf3* mis-expression. In situ hybridisation of otic expression patterns in *Tg(hsp70:fgf3)* embryos following a 30-minute heat shock (HS) at the 10-somite stage (14 hpf). Controls (left-hand panels of each pair of images) were sibling non-transgenic embryos subjected to the same heat shock. Numbers in the dorsal view panels indicate the number of embryos with the phenotype shown and total number (e.g. 13/25) from a mixed batch of transgenic and non-transgenic embryos in each pair of panels; 50% of the batch was expected to be transgenic. **(A-F)** Two hours after heat shock (16 hpf), expression of *hmx3a* expanded to cover the entire otic region (B), but there was only a trace of expression of *hmx2* or *pax5* in the otic placode at this stage. Weak expression of *pax5* in the hindbrain after heat shock (F) did not persist (L). **(G-L)** At 22.5 hpf (8.5 hours after HS), expression of all three genes had now expanded to cover the entire anteroposterior axis of the otic vesicle on the medial side. **(M-R’)** At 36 hpf (22 hours after HS), expression of *hmx3a* remained expanded across the otic anteroposterior axis (N, N’); expression of *hmx2* was strong at the anterior and posterior poles, and weaker in central regions (P, P’), whereas expression of *pax5* resolved into two discrete domains at the anterior and posterior poles of the otic vesicle, and was lost from central regions (R, R’). White arrowheads indicate regions that are normally free of expression in controls; black arrowheads mark ectopic expression in transgenic embryos. A-R are dorsal views showing both otic vesicles, with anterior to the top; M’-R’ are lateral views with anterior to the left. Scale bars, 50 μm (scale bar in A applies to A-L; in M applies to M-R; in M’ applies to M’-R’). For additional examples and time points for *hmx2,* see Fig. 2—Supplemental File 2; for *pax5,* see Fig. 2—Supplemental File 3.

To test whether the milder ear phenotype caused by a later heat shock reflects a failure to establish the early transcriptional responses described above, we also examined expression of anterior markers in *Tg(hsp70:fgf3)* embryos after heat shock at 18 hpf (30 min, 39°C). Unexpectedly, we found that the otic expression of *hmx3a* and *pax5* was very similar to that following an early (14 hpf) heat shock, with a broad band of ectopic expression extending across the entire anteroposterior axis of the otic vesicle by 22.5 hpf (Fig. 1—Supplemental file 2E–H’). This suggests that the loss of competence to generate a complete double-anterior ear after a late heat shock is not due to an inability to express *hmx3a* and *pax5* ectopically throughout the otic epithelium at otic vesicle stages.

### Inhibition of Hh signalling results in a slower and spatially distinct transcriptional response in the otic vesicle

To compare the transcriptional response after *fgf3* mis-expression at 14 hpf with that following conditional Hh pathway inhibition from the same time point, we examined otic expression of anterior marker genes after treatment of wild-type embryos with cyclopamine (100 µM, 14–22.5 hpf; Fig. 3). To confirm the efficacy of cyclopamine treatment, we also examined the expression of *ptch2*, a known target of Hh signalling. Expression of *ptch2* was down-regulated throughout the embryo at 22.5 hpf, but not abolished (Fig. 3—Supplemental file 1). (By contrast, *ptch2* expression is almost entirely lost at 24 hpf in *smo^hi1640Tg/hi1640Tg^* mutants (Chen et al., 2001)). We also checked expression of *etv4* following cyclopamine treatment, but found no major changes in expression at 22.5 hpf (Fig. 3—Supplemental file 1). This result confirmed that there are no strong direct effects of the transient inhibition of the Hh pathway on Fgf signalling activity in the ear, in line with our previous findings after genetic abrogation of Hh signalling (Hammond and Whitfield, 2011).

**Figure 3.**
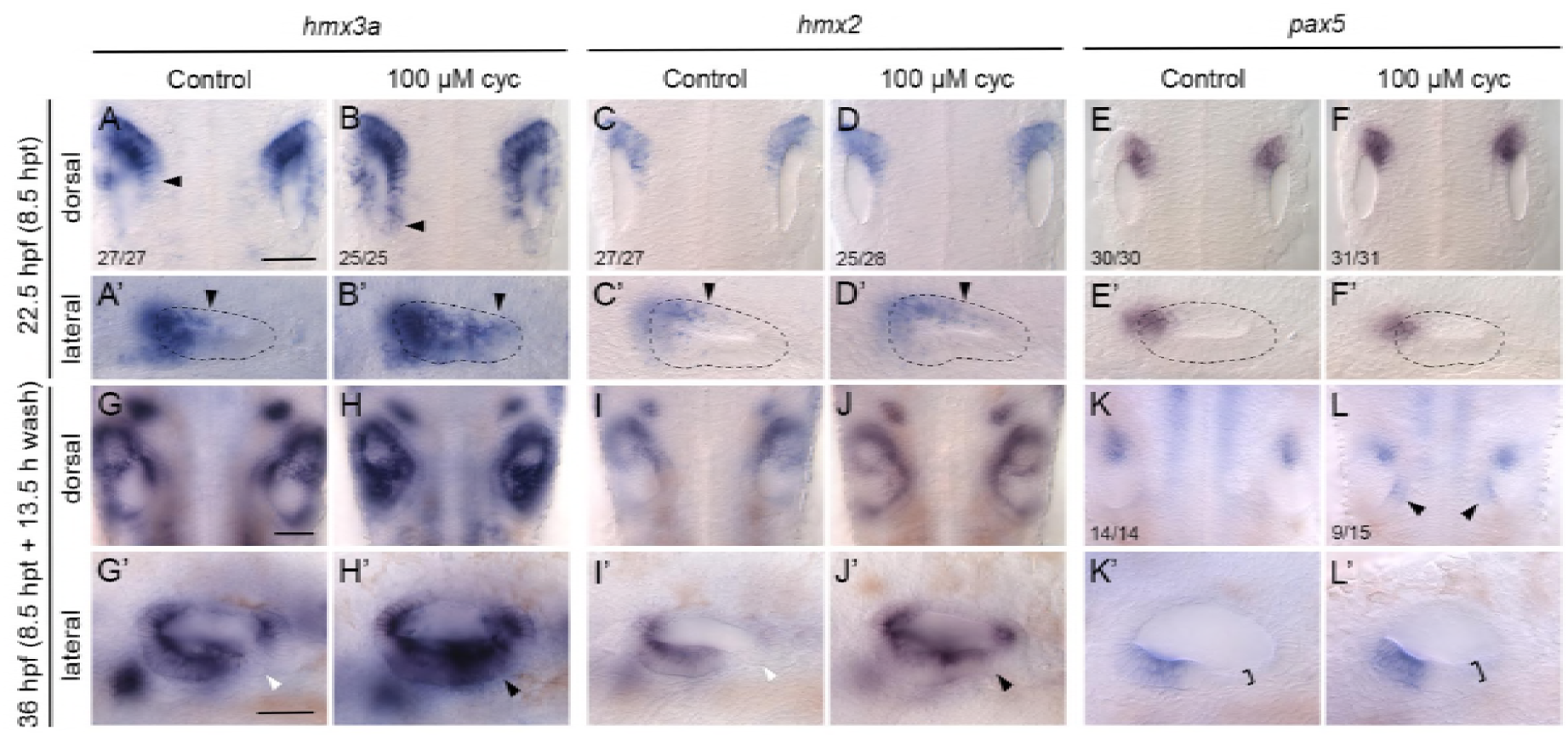
Expression of the otic anterior marker genes *hmx3a, hmx2* and *pax5* after Hh pathway inhibition. Expression of mRNA for anterior otic markers in embryos treated with 100 μM cyclopamine (cyc) from the 10-somite stage (14 hpf) until 22.5 hpf. Controls in the left-hand panels of each pair of images were treated with vehicle (ethanol) only. **(A–F’)** At 22.5 hpf (8.5 hours post initiation of treatment, hpt), expression of *hmx3a* expanded into posterior regions of the otic vesicle (arrowheads); expression of *hmx2* showed a modest expansion and there was no change in the otic expression pattern of *pax5.* Arrowheads in A–D’ indicate posterior extent of otic expression. **(G-L’)** At 36 hpf (8.5 hpt + 13.5 h wash), expression of both *hmx3a* and *hmx2* extended into posteroventral regions of the otic epithelium. White arrowheads indicate regions that are normally free of expression in controls; black arrowheads mark ectopic expression in cyclopamine-treated embryos. By 36 hpf, expression of *pax5* appeared in a new discrete domain in posteromedial otic epithelium after cyclopamine treatment (L, arrowheads); in a lateral view, the epithelium in posterolateral regions was thicker than normal (K’, L’, brackets). A–L are dorsal views showing both otic vesicles, with anterior to the top; A’–L’ are lateral views with anterior to the left. Scale bars, 50 μm (scale bar in A applies to A–F’; in G applies to G–L; in G’ applies to G’–L’).

We examined otic marker genes at two different time points following cyclopamine treatment (22.5 hpf and 36 hpf). Otic expression of both *hmx3a* and *hmx2* was expanded posteriorly on the medial side of the otic vesicle at 22.5 hpf, 8.5 hours after the start of the treatment (Fig. 3A-D’; Fig. 3—Supplemental file 2). Expanded otic *hmx* gene expression was also present by 23 hpf in the *smo^hi1640Tg/hi1640Tg^* mutant, in which the Hh pathway is constitutively inactive (Fig. 3—Supplemental file 2). Importantly, there was no significant difference in the expansion of *hmx3a* expression in the ear at 22.5–23 hpf between the *smo* mutants and cyclopamine-treated embryos, indicating that our cyclopamine treatment regime is effective at suppressing Hh signalling relevant to otic patterning at this stage (Fig. 3—Supplemental file 2).

The spatial pattern of *hmx* expansion in response to Hh inhibition was different to that seen after mis-expression of *fgf3*. Specifically, in cyclopamine-treated embryos or *smo^hi1640Tg/hi1640Tg^* mutants, *hmx3a* and *hmx2* were expressed in a graded fashion across the ear at 22.5 hpf (8.5 hours after treatment), with higher levels anteriorly, rather than in a uniform broad band (compare Fig. 3B with Fig. 2H). To examine later time points, treated embryos were transferred to fresh medium at 22.5 hpf without cyclopamine, as described above. By 36 hpf (13.5 hours post wash), expression of *hmx* genes had expanded further posteriorly to cover most of the ventral floor of the otic vesicle in cyclopamine-treated embryos (Fig. 3G-J’), as observed previously at 30 hpf in *con^tf18b/tf18b^* and *smo^b641/b641^* mutants, both of which have a strong reduction in Hh signalling (Hammond et al., 2003).

Expression of *pax5* was slower to respond following cyclopamine-mediated inhibition of Hh signalling, with no apparent expansion of the expression domain within the otic vesicle at 22.5 hpf (Fig. 3E-F’). These results corroborate our previous observations in *con^tf18b/tf18b^* and *smo^b641/b641^* mutants, where there was little change in the otic expression of *pax5* at 24 hpf (Hammond et al., 2003). However, by the later time point (36 hpf; 13.5 hours post wash), a new, discrete domain of *pax5* expression appeared within posteromedial otic epithelium of cyclopamine-treated embryos (Fig. 3K, L). Anteroposterior asymmetry in treated ears was still evident at this stage: the posterior domain of expression was weaker, and in a more medial position, than the anterior expression domain (Fig. 3K, L). However, the epithelium in posteroventral regions was thicker than normal, indicating development of a duplicate domain of sensory tissue (Fig. 3K’, L’). Taken together, our data indicate that the duplicated anterior domain resulting from either an Fgf gain-of-function or Hh loss-of-function includes a duplication of *pax5* expression, but that otic patterning progresses through completely different intermediate states to achieve this duplicated pattern, depending on the signalling pathway that has been disrupted.

### Expression of Fgf family genes in the otic epithelium following *fgf3* mis-expression or Hh inhibition

Expression of *fgf3* is itself a marker of anterior otic epithelium from 21 hpf (Millimaki et al., 2007), and so can also be used to indicate the presence of a duplicated anterior otic pattern. We therefore examined the expression of *fgf* genes to provide additional confirmation of anterior character in the duplicated ears. To distinguish between expression of the *fgf3* transgene and endogenous *fgf3* expression, we used a probe generated from the *fgf3* 3’ UTR, which is not included in the transgenic construct. In *Tg(hsp70:fgf3)* embryos after early (14 hpf) heat shock, expression of endogenous *fgf3* now appeared in a new domain at the posterior of the otic vesicle at 22.5 hpf (Fig. 4A–B’, arrowheads). Importantly, expression was not found across the entire anteroposterior axis, but was only present at the poles. Expression of endogenous *fgf3* in pharyngeal endoderm beneath the ear was reduced or missing in heat-shocked transgenic embryos (Fig. 4A–B’, asterisks). We also examined the otic expression of *fgf8a* and *fgf10a* (Fig. 4C–F’). These genes are also normally expressed in the anterior of the otic vesicle, but show a less restricted pattern of expression than that of *fgf3,* with weaker expression also normally found in posterior regions (Léger and Brand, 2002; McCarroll and Nechiporuk, 2013; Thisse and Thisse, 2004). Following early heat shock of *Tg(hsp70:fgf3)* embryos, there was little change in the expression of *fgf8a* in the otic epithelium, whereas expression of *fgf10a* was strengthened at both anterior and posterior poles (Fig. 4C–F’).

**Figure 4.**
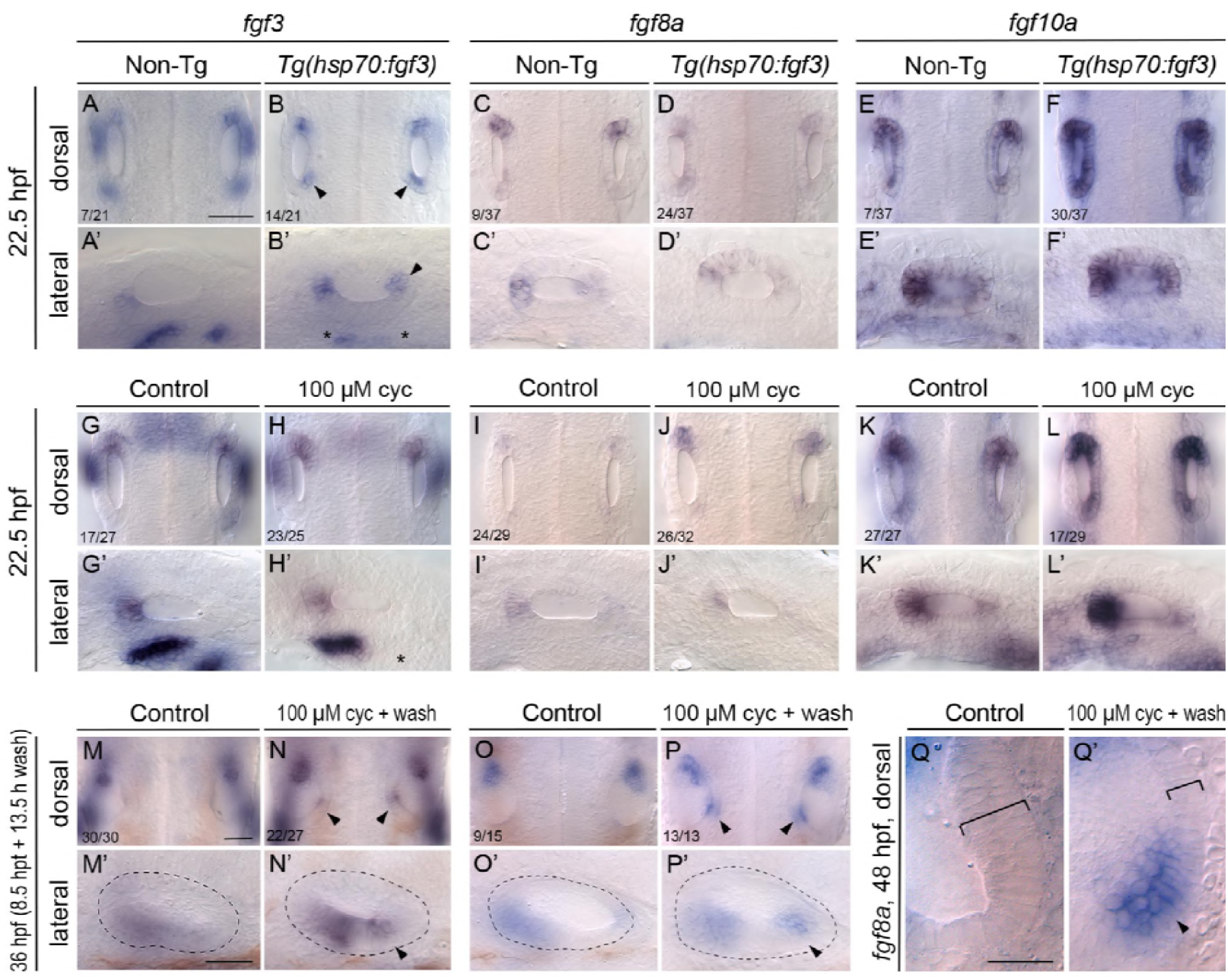
Otic expression of *fgf* genes following mis-expression of *fgf3* or inhibition of Hh signalling. **(A-F’)** In situ hybridisation for otic expression of *fgf* genes *inTg(hsp70:fgf3)* embryos following a 30-minute heat shock (HS) at the 10-somite stage (14 hpf). Controls (left-hand panels of each pair of images) were sibling non-transgenic (Non-Tg) embryos subjected to the same heat shock. Numbers of embryos shown in the dorsal view panels indicate the number showing the phenotype from a mixed batch of transgenic and non-transgenic embryos in each pair of panels; 75% of the batch is expected to be transgenic. A-B’ show staining with a probe specific to the 3’ UTR of *fgf3:* note the ectopic patch of endogenous *fgf3* expression at the posterior otic pole (B, B’; arrowheads) and disruption to *fgf3* expression ventral to the otic vesicle (B’; asterisks) after heat shock in transgenic embryos. Expression of *fgf10a* is strengthened in the otic vesicle of transgenic embryos after heat shock (E-F’). **(G-Q’)** Expression of mRNA for *fgf* genes in embryos treated with 100 μM cyclopamine (cyc) from the 10-somite stage (14 hpf) until 22.5 hpf. Controls in the left-hand panels of each pair of images were treated with vehicle (ethanol) only. Numbers of embryos with the phenotype shown for individual treatments are indicated in the dorsal view panels. There was little change to the otic expression patterns of *fgf3* or *fgf8a* at 22.5 hpf (8.5 hours post treatment) (G-J’), but note the loss of *fgf3* expression ventral to the ear (H’; asterisk). Expression of *fgf10a* in the otic vesicle was strengthened after inhibition of Hh signalling in about 50% of treated embryos (L, L’). At 36 hpf (8.5 hpt + 13.5 h wash), ectopic expression of both *fgf3* and *fgf8a* appeared in a new posteromedial domain in the ears of cyclopamine-treated embryos (M-P’; arrowheads). (Q, Q’) Expression of *fgf8a* in the posterior of the otic vesicle at 48 hpf (8.5 hpt + 25.5 h wash). Ectopic expression has strengthened (arrowhead) and medial epithelium is thinner than normal (brackets). Dorsal views, with anterior to the top. Scale bar in A, 50 μm (applies to A-L); scale bar in A’, 50 μm (applies to A’-L’); scale bar in M, 50 μm (applies to M-P); scale bar in M’, 50 μm (applies to M’-P’); scale bar in Q, 20 μm (applies to Q, Q’).

We also examined the otic expression of *fgf* genes after pharmacological inhibition of Hh signalling. At 22.5 hpf, following cyclopamine treatment from 14 hpf, there was little change in the expression domain or levels of *fgf3* or *fgf8a* in the otic epithelium (Fig. 4G–J’), although there was loss of an *fgf3* expression domain in pharyngeal pouch endoderm ventral to the ear (Fig. 4H’, asterisk). Otic expression of *fgf10a* was strengthened in about 50% of cyclopamine-treated embryos (*n*=17/29) at this early time point, especially at the anterior otic pole (Fig. 4K–L’). At 36 hpf (13.5 hours after cyclopamine wash-out), new discrete domains of *fgf3* and *fgf8a* had appeared at the posterior of the ear, indicating a duplication of anterior otic character (Fig. 4M–P’, arrowheads). By 48 hpf, the duplicated expression domain of *fgf8a* persisted (Fig. 4Q’, arrowhead), and loss of the thickened epithelium characteristic of the posterior macula on the medial wall of the otic vesicle was also apparent (Fig. 4Q, Q’, brackets). These data demonstrate that in both Fgf gain-of-function and Hh loss-of-function contexts, the duplicated anterior otic character includes expression of *fgf* genes.

### Loss of *hmx3a* function results in a fusion of sensory maculae and otoliths, and a reduction in anterior otic character

Given the early anterior-specific otic expression of *hmx3a* (Feng and Xu, 2010), the dependence of this expression on Fgf signalling (Adamska et al., 2000; Hammond and Whitfield, 2011; Kwak et al., 2002), and the rapid change in otic *hmx3a* expression after mis-expression of *fgf3* or Hh inhibition (this work), we hypothesised that *hmx3a* is required for normal otic anterior development. A previous study using morpholino-mediated knockdown suggested a requirement for both *hmx3a* and *hmx2* in acquisition of anterior otic identity and expression of *pax5* (Feng and Xu, 2010). However, the effects of individual gene knockdown or mutation were not reported. To test the individual requirement for *hmx3a* function in the acquisition of otic anterior identity, we examined the ear phenotype in homozygous mutants for a recessive truncating allele lacking the homeodomain, *hmx3a^SU3^*, which we generated using CRISPR/Cas9 technology (Fig. 5A; Materials and Methods). In homozygous *hmx3a^SU3/SU3^* mutants, the otoliths were positioned close together at 33 hpf, were side by side at 48 hpf, had started to fuse at 66 hpf and had fully fused by 4 dpf (Fig. 5B, C and data not shown). This phenotype appeared to be fully penetrant (38/143 embryos from a cross between heterozygous parents; 26.6%). Semicircular canal pillars and the dorsolateral septum were present in the ears of mutant embryos, although formation of the ventral pillar was delayed. Overall, the ear shape appeared more symmetrical than in wild-type siblings (Fig. 5B, C). We imaged ears from three mutant embryos at 3 dpf to analyse sensory patch formation (Fig. 5D–E’ and Fig. 5—Supplemental file 1). In all three ears imaged, the two maculae appeared fused or closely juxtaposed. Although the anterior and posterior elements of the fused macula were not obviously distinct, the overall shape retained some anteroposterior asymmetry. Hair cells of the anterior (utricular) macula were displaced medially, and in one of three ears imaged, were reduced in number. In the two other examples, however, normal numbers of hair cells were present (Fig. 5—Supplemental file 1). The posterior macula was misshapen, and lacked the anterior extension present in the wild type. All three cristae were present (*n*=3 ears; Fig. 5D’, E’ and Fig. 5—Supplemental file 1).

**Figure 5.**
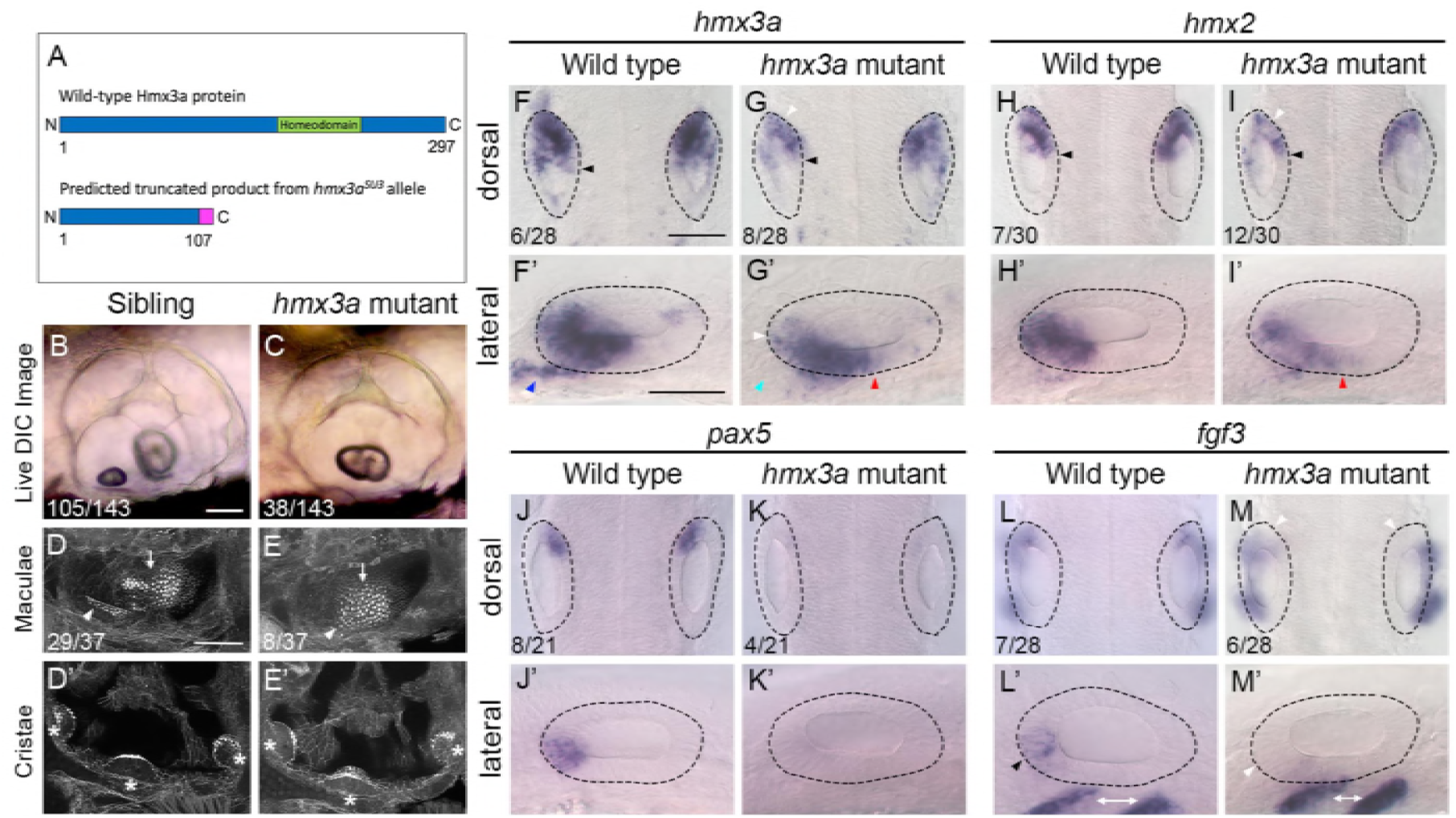
Fused otoliths and sensory maculae, and reduction of anterior otic character, in *hmx3a^SU3/SU3^* mutants. **(A)** Schematic diagram showing the predicted truncated product for the *hmx3a* allele. The mutation was generated using a CRISPR sgRNA targeting sequence in exon 2 upstream of the DNA-binding homeodomain (green). The predicted truncated protein produced by the *hmx3a^SU3/Sü3^* allele contains a Thr to Gly substitution at amino acid 107, followed by a stretch of 10 further incorrect amino acids (magenta). The truncated protein lacks the homeodomain. **(B, C)** Differential interference contrast (DIC) images of ears in live embryos at 3 dpf (72 hpf). Numbers of embryos in a batch from a mating between heterozygous parents are given. Note the fused otolith in the *hmx3a^SU3/SU3^* mutant ear (C). **(D–E’)** FITc-phalloidin stains of the sensory maculae (D, E) and cristae (D’, E’) in the ear at 3 dpf (72 hpf). Numbers of embryos showing the phenotype from a cross between heterozygous parents are shown. White arrowhead: anterior macula; white arrow: posterior macula; asterisks indicate cristae. Additional examples are shown in Fig. 5—Supplemental file 1. **(F–M’)** In situ hybridisation for otic anterior markers at 24 hpf in genotyped wild-type and *hmx3a^SU3/S 3^* mutant embryos. The dotted outline marks the outer edge of the otic epithelium. Black arrowheads in F–I indicate the extent of *hmx* expression in medial epithelium; white arrowheads indicate areas of reduced expression levels; blue arrowhead in F’ marks presumed otic or anterior lateral line neuroblasts; light blue arrowhead in G’ indicates loss of expression in this area; red arrowheads in G’, I’ mark expansion of expression in ventral otic epithelium. Black arrowhead in L’ indicates anterior otic expression domain of *fgf3,* lost in M, M’ (white arrowheads); white double-headed arrows mark expression of *fgf3* in pharyngeal pouch endoderm. Numbers in panels F–M indicate numbers of embryos genotyped as either wild type or homozygous mutant that showed the representative expression patterns illustrated. Scale bars, 50 μm (scale bar in B applies to B, C; scale bar in D applies to D–E’; scale bar in F applies to F–I, JM; scale bar in F’ applies to F’–I’, J’–M’).

To understand the basis of the *hmx3a^SU3/SU3^* mutant otic phenotype at 3–4 dpf, we examined expression of markers at earlier (otic vesicle) stages. At 24 hpf, expression of both *hmx3a* and *hmx2* was reduced in intensity within the otic epithelium. On the medial side of the ear, the spatial extent of *hmx3a* and *hmx2* expression was unchanged (Fig. 5F–I, black arrowheads), but levels were reduced (white arrowheads); anteroventrally, there was a reduction in *hmx3a* expression in presumed neuroblasts (Fig. 5F’, G’, blue and light blue arrowheads), and a mild posterior expansion of the spatial extent of expression for both genes in ventral otic epithelium (Fig. 5G’, I’, red arrowheads). Expression of the anterior markers *pax5* and *fgf3* was drastically reduced within anterior otic epithelium in *hmx3a^SU3/SU3^* mutants at 24 hpf (Fig. 5J–M’, arrowheads), although expression of *fgf3* in pharyngeal pouch endoderm ventral to the ear was unaffected (Fig. 5M, M’, double-headed white arrows). Expression of the same markers in *hmx3a^SU3/SU3^* mutants at 27 hpf was similar, but otic expression of *hmx2* was more strongly reduced than that of *hmx3a*, especially in the anterior pole in the area corresponding to the normal expression domain of *fgf3* and *pax5* (Fig. 5—Supplemental file 2). Expression of the posterior marker *fsta* at 30 hpf did not reveal any significant duplication of expression in anterior otic epithelium in *hmx3a^SU3/SU3^* mutant ears (Fig. 5—Supplemental file 2). Taken together, the results suggest that a loss of *hmx3a* function results in a similar phenotype to that of *fgf3^-/-^* (*lia^t21142/t21142^*) mutants (Hammond and Whitfield, 2011; Kwak et al., 2006; Maier and Whitfield, 2014). Although some anteroposterior asymmetry has been lost, the phenotype is not as strong as the double-posterior duplications that result from inhibition of all Fgf signalling or over-activity of Hh signalling, which show a complete loss of the anterior macula and lateral crista, and duplication of elements of the posterior macula (Hammond et al., 2010; Hammond and Whitfield, 2011). We conclude that *hmx3a* function is required for normal anterior otic expression of *pax5* and *fgf3*. However, loss of *hmx3a* function is not sufficient to result in a complete loss of anterior character and duplication of posterior structures at the anterior of the ear.

### Mis-expression of *hmx3a* is not sufficient to result in a duplication of anterior otic identity

As otic expression of the anterior markers *pax5* and *fgf3* is strongly reduced in *hmx3a^SU3/SU3^* single mutants, and because expression of *hmx3a* is an early transcriptional response to manipulations of both Fgf and Hh signalling, we hypothesised that mis-expression of *hmx3a* alone would be sufficient to drive the expression of *pax5* and *fgf3* in the posterior of the otic placode and to give rise to a double-anterior duplication, bypassing the requirement for Fgf or Hh pathway manipulation. To test this idea, we created a transgenic line driving expression of the *hmx3a* coding sequence under the control of the *hsp70* heat-shock promoter. A 60-minute heat shock of *Tg(hsp70:hmx3a)* embryos at 12 hpf resulted in a robust and widespread expression of the *hmx3a* transgene two hours later (Fig. 6A–D). To avoid any disruption of otic placode induction, and to be comparable to the *fgf3* heat-shock experiments, we heat-shocked *Tg(hsp70:hmx3a)* embryos at 14 hpf to induce systemic mis-expression of *hmx3a*. After 30 minutes at 39°C, heat-shocked embryos were incubated at 33°C for 30 minutes before being returned to 28.5°C and incubated until 22.5 hpf, when they were fixed for processing and analysis, or until 3 dpf for assessment of any ear duplication (Fig. 6E–Q’).

**Figure 6.**
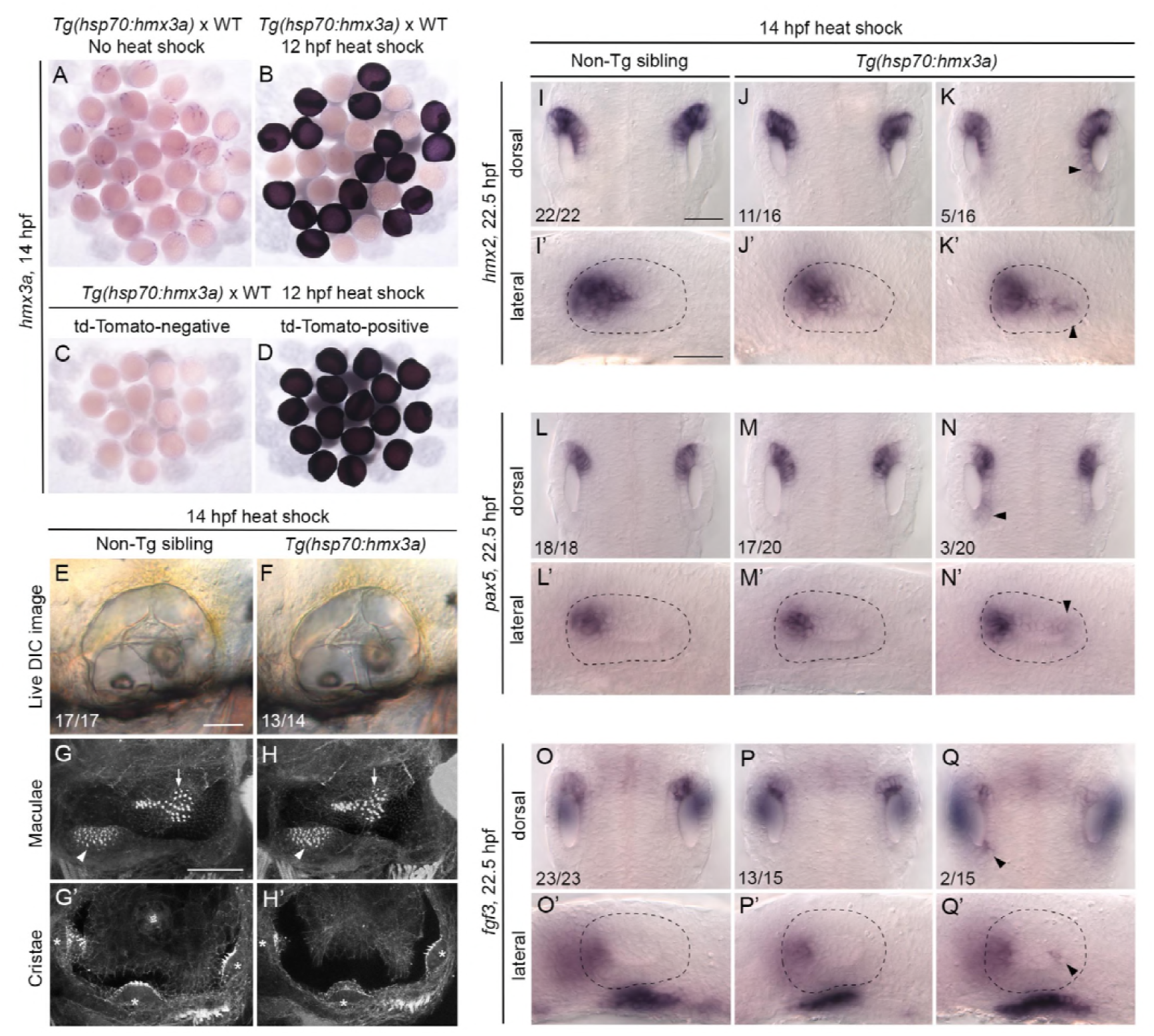
Mis-expression of *hmx3a* is not sufficient to generate an anterior ear duplication. **(A-D)** Control experiments to check for successful expression of the *hmx3a* transgene after heat shock at 12 hpf. Embryos were fixed and stained by in situ hybridisation two hours later, at 14 hpf. **(A, B)** A mixed batch of embryos from a cross between a fish hemizygous for the transgene and a wild type (WT). All embryos (31/31) showed the normal pattern of expression of *hmx3a* in the absence of heat shock (A). After a 60-minute heat shock at 12 hpf, ∼50% of the batch (17/30) showed strong, systemic expression of the transgene at 14 hpf, as expected (B). All embryos shown in B were stained in the same tube. **(C, D)** Embryos heat-shocked for 60 minutes at 12 hpf were sorted on the basis of tdTomato expression before fixing. All tdTomato-negative embryos (14/14) were also negative for expression of the *hmx3a* transgene (C); all tdTomato-positive embryos (16/16) were also positive for *hmx3a* transgene expression (D). **(E, F)** Live DIC images of ears of non-transgenic (E) and transgenic (F) sibling embryos at 3 dpf (72 hpf), after a 30-minute heat shock at 14 hpf. **(G-H’)** Confocal images of FITC-phalloidin-stained ears at 3 dpf (72 hpf). Position and size of the two maculae (G, H, arrowheads and arrows) and three cristae (G’H’, asterisks) were normal in both non-transgenic and transgenic sibling embryos after heat shock. **(I-Q’)** In situ hybridisation for otic marker genes in non-transgenic and *Tg(hsp70:hmx3a)* sibling embryos after a 30-minute heat shock at 14 hpf. Dorsal views of both otic vesicles (I-Q) and lateral views of a single otic vesicle (I’-Q’) are shown. Note weak ectopic expression of *hmx2* and *pax5,* and posterior otic expression of *fgf3,* in the otic vesicles of a minority of transgenic embryos (right hand column; arrowheads). Numbers of embryos showing the phenotypes are shown for each panel. WT, wild type (AB strain). Scale bar in E, 50 μm (applies to E–F); scale bar in G, 50 μm (applies to G–H’); scale bar in I, 50 μm (applies to I–Q); scale bar in I’, 50 μm (applies to I’–Q’).

Despite robust expression of the *hmx3a* transgene, the ears of *Tg(hsp70:hmx3a)* embryos heat-shocked for 30 minutes at 14 hpf did not recapitulate the duplicated double-anterior otic phenotype seen in *Tg(hsp70:fgf3)* embryos. Position and number of the otoliths, morphology of the semicircular canal pillars and position of the sensory patches in heat-shocked embryos were normal at 3 dpf (Fig. 6E–H’; compare with Fig. 1B). At 22.5 hpf, otic vesicles were slightly smaller and rounder in heat-shocked transgenic embryos than those in heat-shocked non-transgenic siblings, but markers were expressed normally in most cases (Fig. 6I–Q’). Otic expression of *hmx2* was mildly up-regulated in a graded fashion (higher at the anterior) in 5/16 transgenic embryos at 22.5 hpf (Fig. 6I–K’), similar to the de-repression of *hmx* expression seen after Hh inhibition in wild-type embryos. There was also a mild up-regulation of *pax5* expression in posterior otic epithelium at 22.5 hpf in 3/20 embryos (Fig. 6L–N’), but *pax5* was never expressed in a broad zone as in the *Tg(hsp70:fgf3)* embryos. A weak patch of *fgf3* expression appeared in posterior otic epithelium at 22.5 hpf, similar to the duplicated zone of endogenous *fgf3* expression in *Tg(hsp70:fgf3)* embryos, but in only 2/15 embryos (Fig. 6O–Q’).

To check that the *hmx3a* transgene was functional, we sequenced it from genomic DNA of transgenic embryos, which indicated that the open reading frame was intact (data not shown). We also examined the phenotype of transgenic embryos after an even earlier heat shock, during otic placode induction (8–9 hpf and 10–11 hpf). Here, we saw a range of otic abnormalities in 80% of transgenic embryos (*n*=106), including missing otoliths, but some embryos also had small heads and eyes (Fig. 6—Supplemental file 1). Ear patterning appeared normal in about 20% of transgenic embryos heat-shocked at these earlier stages. Longer (1- or 2-hour) heat shocks at 14–15 hpf also resulted in normal otic patterning (*n*=49; Fig. 6—Supplemental file 1). We conclude that the *hmx3a* transgene is likely to be functional, but that its mis-expression alone during otic placode stages (14–15 hpf, which should result in strong systemic expression until at least 17 hpf) cannot substitute for Fgf mis-expression or Hh inhibition in the generation of a double-anterior duplicated ear. Up-regulation of *hmx3a* in the ear at later stages, beyond 18 hpf, was not sufficient either, as our late *fgf3* heat shock experiments demonstrated (Fig. 1—Supplemental File 2).

### A dynamical model of anteroposterior patterning in the zebrafish ear

Taken together, our data and those from previously-published studies suggest a temporal hierarchy of events for otic anteroposterior patterning dependent on extrinsic sources of Fgf and Hh signalling (Fig. 7). To assess whether this network of inferred genetic regulatory interactions can account for the dynamic expression patterns we observe, we developed a mathematical model of otic anteroposterior patterning in the wild-type ear and following manipulation of the Fgf and Hh signalling pathways. The model is based on a set of differential equations describing the genetic interactions in the otic epithelium outlined in Figure 7A. In addition, patterning in the model is dependent on the existence of two sources of spatial information. First, we assume that otic competence to express *fgf* genes in response to Fgf and Hmx3a protein is localised to the two poles of the developing otic vesicle. This is necessary in the model to ensure that induced endogenous *fgf* mRNA expression in the otic epithelium (*fgf_i_*) is restricted to the poles, even when *fgf3* is expressed uniformly throughout the tissue following heat shock. Second, we represent the effect of *fgf* mRNA expression in rhombomere 4 as an anterior-to-posterior gradient of extrinsic Fgf (Fgf_e_) protein, present at high levels up to 30% of the otic vesicle length (corresponding to the position of the rhombomere 4/5 boundary), and forming a decreasing spatial gradient across the remainder of the otic axis (Fig. 7A and Fig. 8—Supplemental File 1). Although we do not have a measure of actual Fgf protein concentration, our assumption is supported by measurements of fluorescence across the otic anteroposterior axis in the *Tg(dusp6:d2EGFP)* reporter line, which expresses a destabilised GFP variant as an indirect readout of Fgf activity (Molina et al., 2007) (Fig. 7—Supplemental file 1).

**Figure 7.**
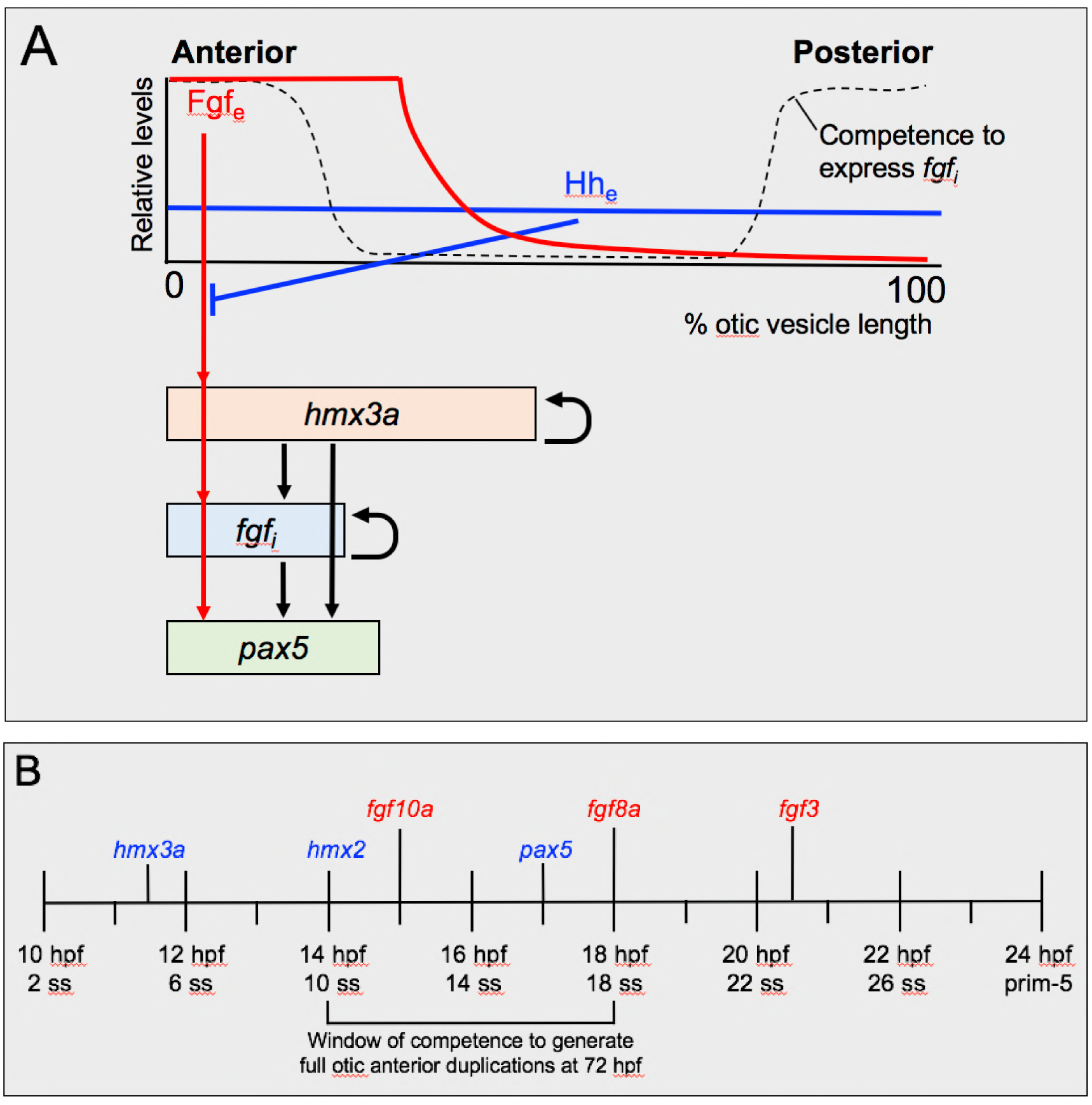
Proposed gene network and timeline for the acquisition of anterior identity in the zebrafish otic vesicle. **(A)** Proposed gene regulatory network. **(B)** Schematic timeline showing sequential onset of expression of anterior markers in the zebrafish otic placode and vesicle. Diagrams in A and B are based on the results of this study, together with previously published data (Feng and Xu, 2010; Hammond et al., 2003; Hammond et al., 2010; Hammond and Whitfield, 2011; Kwak et al., 2006; Léger and Brand, 2002; Maier and Whitfield, 2014; McCarroll and Nechiporuk, 2013; Millimaki et al., 2007). Abbreviations: Fgf_e_, extrinsic Fgf protein; fgf, intrinsic (otic vesicle) *fgf* gene expression; Hh_e_, extrinsic Hedgehog protein; hpf, hours post fertilisation; ss, somite stage.

We assume a spatially uniform level of Hh signalling throughout the otic epithelium (see discussion in (Hammond et al., 2003), and that Hh signalling antagonises the effects of Fgf signalling on otic anterior marker genes (*fgf_i_*, *hmx3a* and *pax5*) by increasing their response threshold for Fgf-induced expression. This functional attenuation is unlikely to be at the level of an immediate target of Fgf signalling such as *etv4*, as Hh inhibition did not result in major changes to *etv4* expression ((Hammond and Whitfield, 2011); this work). One possibility is that it could reflect integration of activity of the two signalling pathways at the level of binding sites in the promoters of the otic anterior genes. In addition, we propose that Hmx3a, together with Fgf and Hh, regulates its own expression and that of other genes in the network. Currently, our data do not distinguish whether these regulatory relationships are direct or indirect.

The dynamic behaviour of the model is presented in Figure 8 (for full details, see Fig. 8—Supplemental File 1). In wild-type embryos (Fig. 8, left-hand column), expression of *hmx3a* and *pax5* is triggered in anterior otic tissue. The extent of expression is determined by the spatial reach of the extrinsic Fgf protein (Fgf_e_) gradient from rhombomere 4. Although all cells in the model are competent to express the anterior markers *hmx3a* and *pax5*, they do not receive sufficient Fgf_e_ to do so at the posterior otic pole in a wild-type embryo. After transient heat shock-induced systemic mis-expression of *fgf3* at 14 hpf (Fig. 8, middle column), expression of both *hmx3a* and *pax5* is induced across the entire anteroposterior axis. However, the ability of heat shock-induced *fgf3* mis-expression to trigger endogenous intrinsic *fgf* (*fgf_i_*) expression requires the coincidence of both Fgf protein and competence to express *fgf_i_* at the poles, and so *fgf_i_* is not induced in the middle of the otic axis. After decay of heat shock-induced Fgf protein, expression of *pax5* is lost from central regions, but is maintained at the poles by Fgf signalling from *fgf_i_* expression. By contrast, expression of *hmx3a* is maintained in central regions due to its autoregulation. De-repression of anterior markers after Hh pathway inhibition (Fig. 8, right-hand column) results from a lowering of the threshold for response to Fgf signalling, establishing duplicate expression domains of *pax5* and *fgf3* at the posterior pole. Thus, although both heat shock-driven mis-expression of *fgf3* and inhibition of Hh signalling result in anterior duplications (Fig. 8; compare the patterns at the 36 hpf time point), the transient dynamics exhibited by the model at earlier time points are distinct.

**Figure 8.**
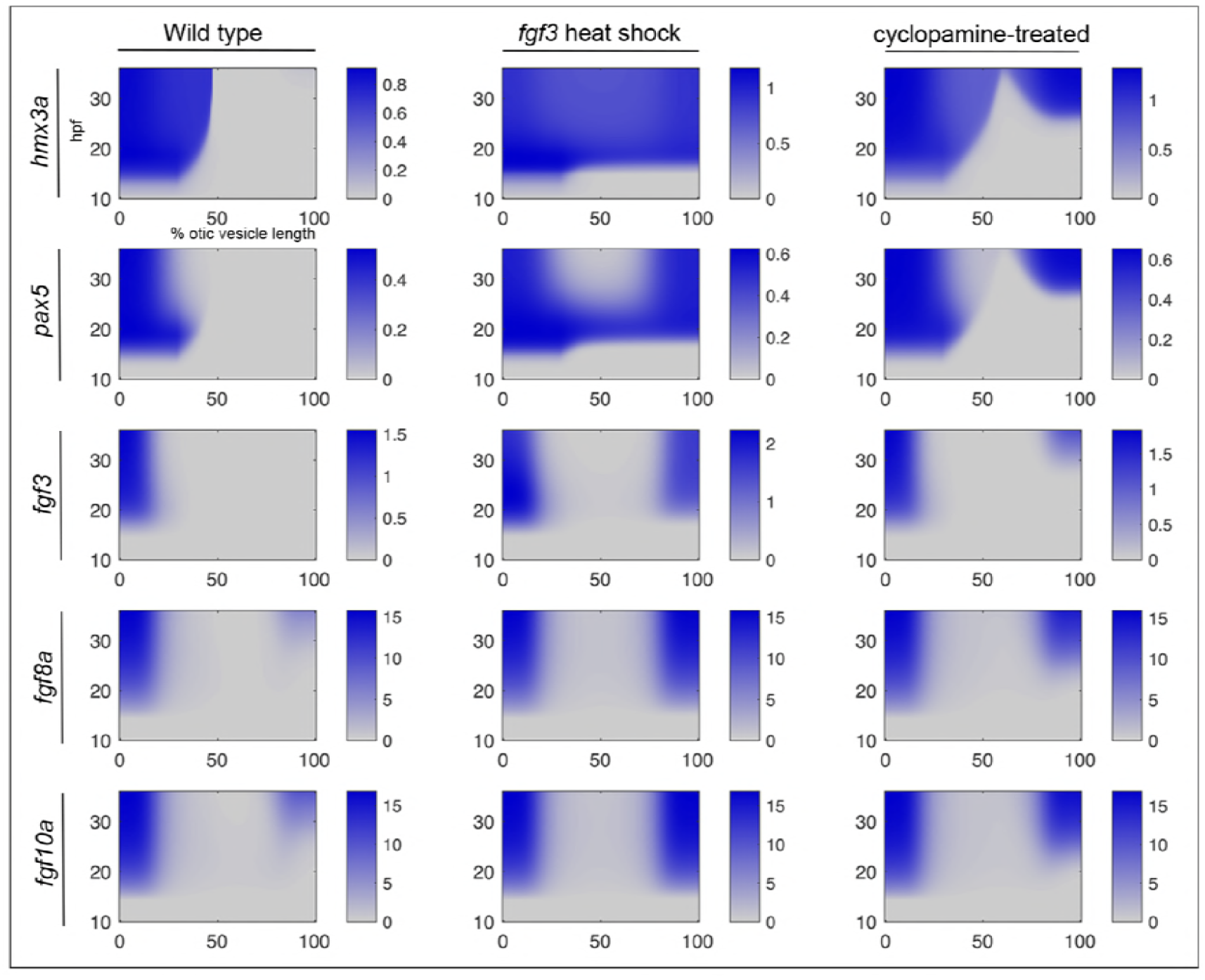
A model for the acquisition of anterior identity in the zebrafish otic vesicle. Solutions of a differential equation-based model to describe gene expression dynamics of the proposed network. Endogenous mRNA expression levels (blue) are shown as a function of position along the otic anteroposterior axis (*x* axis; % otic vesicle length from the anterior end) and time (*y* axis; hpf). Levels are shown arbitrary units in the bars to the right of each panel. Exogenous *fgf3* mRNA from the transgene after heat shock is not shown. Left-hand column: wild type; middle column: with transient heat-shock induction of *fgf3* at 14 hpf; right-hand column: with inhibition of Hh signalling (cyclopamine treatment from 14–22.5 hpf). For full details, see Figure 8—Supplemental file 1.

## Discussion

### Different transcriptional dynamics in the otic vesicle in response to manipulations of Fgf and Hh signalling

The zebrafish otic placode is a convenient system in which to understand the gene network dynamics that lead to asymmetries along the axis of a developing organ. Asymmetries in gene expression are evident from early (otic placode) stages, but the system is clearly equipotential, since either a gain of Fgf signalling or a loss of Hh pathway activity at otic placode stages can produce remarkably similar double-anterior zebrafish ears at 3 dpf (Hammond et al., 2003; Hammond and Whitfield, 2011); this work. Interestingly, we have shown here that this final duplicated pattern arises via very different intermediate states in terms of gene expression patterns, depending on the signalling pathway that has been disrupted. Mis-expression of *fgf3* at 14 hpf leads to a rapid loss of asymmetry, with broad expansion of anterior otic markers across the entire anteroposterior axis of the ear within a few hours of heat shock-driven mis-expression. Expression of *pax5*, which is required for normal development of the anterior (utricular) macula (Kwak et al., 2006), later resolves into two discrete domains. By contrast, initial asymmetries in gene expression persist for several hours after inhibition of Hh pathway activity, with new duplicate expression domains of anterior markers (*pax5, fgf3* and *fgf8a*) only appearing nearly a day later at the posterior otic pole. We have identified *hmx3a* as an early otic transcriptional response to manipulations of both signalling pathways. However, although a loss of *hmx3a* demonstrates its requirement for normal otic patterning, this does not result in a complete double-posterior duplication, and mis-expression of *hmx3a* does not appear to be sufficient to drive the formation of a double-anterior ear.

Our data and mathematical model suggest that the Fgf/Hh system is sufficient to pattern the anteroposterior axis of the ear. In our scheme, there is only one input (extrinsic Fgf activity) that has a graded distribution across the otic anteroposterior axis. Notably, there is no need to infer an opposing graded input of extrinsic signalling activity that is high at the posterior of the ear. Although Retinoic Acid (RA) is thought to form such a gradient, and contributes to anteroposterior patterning in both the chick and zebrafish ear (Bok et al., 2011; Radosevic et al., 2011), its activity can clearly be over-ridden by manipulations of Fgf or Hh signalling in generating either double-anterior or double-posterior zebrafish ears. Our model therefore differs from other models of axial patterning, for example in generation of dorsoventral pattern in the vertebrate neural tube. Here, information from two anti-parallel noisy gradients is integrated and refined by cross-repressing interactions between target genes, providing precise positional information along the axis (Briscoe and Small, 2015; Zagorski et al., 2017). However, the sufficiency of our network and model does not necessarily rule out a contribution from the RA gradient in generating correct anteroposterior patterning in the wild-type ear.

At present, we do not have a full mechanistic explanation for the differences in response dynamics after manipulations of the Fgf and Hh signalling pathways. Although *hmx3a* responds rapidly to manipulation of Fgf signalling, its regulation may well be indirect; a recent study identified only one gene, *Etv5*, as a direct up-regulated target of Fgf signalling during induction of otic-epibranchial precursor cells in the chick (Anwar et al., 2017). In zebrafish, transcription of *etv4* and *spry4* is known to be an early response to Fgf signalling (Raible and Brand, 2001; Roehl and Nüsslein-Volhard, 2001; Scholpp and Brand, 2004), with *spry4* expression appearing within one hour of implantation of a bead coated with Fgf8 protein during epiboly stages (Scholpp and Brand, 2004). Our work here shows that robust, systemic expression of *etv4* occurs within two hours of the onset of heat shock in *Tg(hsp70:fgf3)* embryos; we had previously shown strong expression of *etv4* in the otic placode four hours after heat shock (Hammond and Whitfield, 2011). Thus, Etv4 is a good candidate for an immediate early transcriptional effector of Fgf signalling in our proposed genetic network. However, as *etv4* mRNA expression is not strongly perturbed by Hh pathway inhibition ((Hammond and Whitfield, 2011); this work), effects of Hh and Fgf on *hmx3a* expression are likely to be integrated further downstream, for example at the level of the *hmx3a* promoter. The slower response of *hmx3a* transcription to Hh inhibition might reflect the persistence of Hh pathway effectors, such as Gli activator proteins, which must be degraded before the effect of inhibiting Smoothened with cyclopamine can take effect.

As the otic vesicle develops, additional levels of regulation are likely to contribute to the regulation of *hmx3a* and other genes in the network. For example, in the chick, regulation of *Hmx3* expression in the dorsolateral otocyst has recently been shown to be influenced by both Shh and non-canonical BMP signalling through PKA and GLI3R (Ohta et al., 2016). In addition, negative feedback on otic *fgf* expression via *sprouty* genes (Léger and Brand, 2002) is likely to help to restrict gene expression to the poles and sharpen expression domain boundaries within the otic epithelium.

### Requirement for intrinsic factors at the otic anterior and posterior poles in establishing a duplicate pattern after signalling pathway manipulation

One of the intriguing features of the double-anterior ears is that systemic mis-expression of an anteriorising factor (*fgf3*) gives rise to two defined and separate anterior maculae with mirror-image symmetry, rather than establishing uniform anterior identity across the entire medial otic domain. The final duplicate pattern develops despite the initial broad expression of anterior markers after heat shock, which demonstrates that the entire medial side of the otic placode and vesicle is competent to express *hmx* genes and *pax5* in response to Fgf signalling. However, a day after heat shock, expression of *pax5* is lost from the centre of this domain and only maintained at the anterior and posterior poles of the otic vesicle, suggesting either that expression is subsequently repressed in the central domain, or that an intrinsic factor or factors is required to maintain expression at the poles. Attractive candidates for the latter role include *atoh1a*, which is expressed in discrete domains at the anterior and posterior otic poles at 14 hpf (Millimaki et al., 2007). Atoh1a is thought to act in a positive feedback loop together with Fgf signalling in the zebrafish ear (Millimaki et al., 2007; Sweet et al., 2011). Fgf pathway activity is also observed at both poles of the otic vesicle at 24 hpf (this work), 28 hpf and 50 hpf using a destabilised fluorescent transgenic reporter, *Tg(dusp6:d2EGFP)* (Molina et al., 2007). We hypothesise that a positive feedback loop involving a pole-specific factor and all three *fgf* genes contributes to the maintenance of anterior-specific gene expression and generation of the double-anterior pattern. This builds on previous feedback models for anterior otic patterning and the regulation of otic *pax5* expression (Feng and Xu, 2010; Kwak et al., 2006).

A similar broad medial expansion of *hmx3a* and *pax5* has been recently reported to result from systemic mis-expression of *sox2* or *sox3* at 12.5 hpf (Gou et al., 2018). However, this early mis-expression results in a smaller and mis-shapen otic vesicle (most likely due to a disruption of otic induction), and phenotypes were not followed beyond 30 hpf. It will be interesting to see whether a duplicated anterior pattern results from these manipulations.

### Comparison of the *hmx3a^SU3/SU3^* and *fgf3^t21142/t21142^* mutant otic phenotypes in the zebrafish

The otic phenotype of *hmx3a^SU3/SU3^* single mutants closely resembles that of *hmx3a/hmx2* double morphants (Feng and Xu, 2010), and of *fgf3^t21142/t21142^* mutants (Hammond and Whitfield, 2011; Kwak et al., 2006; Maier and Whitfield, 2014). The similarity of the *fgf3* and *hmx3a* otic mutant phenotypes suggests that a major role for the extrinsic Fgf3 signal is to activate *hmx3a* expression in anterior otic epithelium. Note that pharmacological inhibition of all Fgf signalling (Hammond and Whitfield, 2011) or over-activity of the Hh pathway (Hammond et al., 2010) both result in a stronger otic phenotype than in *hmx3a^SU3/SU3^* mutants. The retention of some anteroposterior asymmetries in gene expression and the fused sensory macula in *hmx3a^SU3/SU3^* mutants, together with the presence of the lateral crista, suggest that the *hmx3a^SU3/SU3^* otic phenotype, like that of *fgf3^t21142/t21142^* mutants, does not represent a complete double-posterior duplication. We also failed to see strong ectopic expression of the posterior marker *fsta* at the anterior of the ear in *hmx3a^SU3/SU3^* mutants, although this is a less reliable indicator of posterior duplication; it is expressed at both poles of the ear following strong Fgf inhibition (Hammond and Whitfield, 2011), but lost altogether in the extreme double-posterior ears that can result from elevated Hh signalling (Hammond et al., 2010).

Despite the similarities between the loss-of-function phenotypes for *fgf3* and *hmx3a* in the zebrafish ear, the gain-of-function effects for each of the two genes are strikingly different. Whereas mis-expression of *fgf3* at 14 hpf reliably generates a complete double-anterior ear, mis-expression of *hmx3a* at the same time point had very little effect on otic development. It is remarkable just how robust the embryo is to this kind of perturbation, considering that the systemic high levels of transgene expression must be energetically expensive to support. Indeed, there is usually some transient developmental delay after heat shock, but gross patterning of the ear at 3 dpf appeared normal in *Tg(hsp70:hmx3a)* heat-shocked embryos.

Why, then, is *hmx3a* ineffective in establishing duplicate anterior development when mis-expressed? It is possible that it needs to be delivered together with *hmx2*; the two genes are tightly linked on zebrafish chromosome 17 (Wotton et al., 2009), spatially co-expressed in the zebrafish otic vesicle (although with different temporal onset) (Feng and Xu, 2010), and are known to have partially overlapping roles in the mouse ear (Wang et al., 2004). A predicted *hmx3b* gene (RefSeq XM_017358610.2) is also present in the zebrafish genome on chromosome 12, although it does not appear to be expressed in the ear (S. England and K. Lewis, unpublished). If a second Hmx family protein or other binding partner was limiting, this might explain the lack of activity of the mis-expressed *hmx3a* transcript. Alternatively, Hmx3a could act as a competence factor, only functioning in the context of high Fgf or low Hh signalling to initiate duplicate anterior otic development. It is also possible that Fgf signalling abrogates an unidentified negative regulator of the otic anterior gene network at the same time as activating the expression of *hmx3a*. In the presence of such an inhibitor, systemic over-expression of *hmx3a* would be ineffective at activating the expression of genes such as *hmx2, pax5* and *fgf3* in posterior otic domains.

### Comparison of the effects of loss of *Hmx3* function on otic development between zebrafish and amniotes

Anterior-specific otic expression of *Hmx3* and *Hmx2*, including their temporal order of expression onset in the ear, is conserved between zebrafish, mouse and chick (Feng and Xu, 2010; Herbrand et al., 1998; Rinkwitz-Brandt et al., 1995; Wang et al., 1998). Loss of *Hmx3* function in the mouse causes a range of reported otic defects with variable penetrance and expressivity, which depend on the nature of the targeted mutant allele. A homozygous targeted deletion of exon 1 and part of exon 2 of *Hmx3* resulted in a variable disruption of the lateral (horizontal) and posterior semicircular canal ducts, and loss of the lateral (horizontal) crista (Hadrys et al., 1998). A weaker phenotype was seen after disruption of the homeodomain in exon 3 of *Hmx3*; in these mutants, all three semicircular canal ducts were present, but the lateral (horizontal) ampulla and crista were missing. The utricular and saccular maculae were juxtaposed in a common utriculosaccular chamber (Wang et al., 2004; Wang et al., 1998), as we found in the zebrafish *hmx3a^SU3/SU3^* mutant. A notable difference between the mouse and zebrafish mutants is the presence of all three cristae, including the lateral crista, in the zebrafish *hmx3a^SU3/SU3^* mutants. Formation of the ventral pillar for the lateral canal was also present, although delayed. It will be interesting to see whether mutations in *hmx2* (not currently available) affect morphogenesis of the zebrafish semicircular canal system; in the mouse, targeted disruption of *Hmx2* results in a loss of all three semicircular canal ducts, with partial or complete loss of some ampullae and cristae, in addition to a fused utriculosaccular chamber (Wang et al., 2001). In humans, *HMX3* and *HMX2* are located together, close to *FGFR4,* on chromosome 10; hemizygous microdeletions that remove all three genes are thought to be causative for syndromes characterised by inner ear morphological anomalies, vestibular dysfunction and sensorineural hearing loss (Miller et al., 2009; Sangu et al., 2016).

In conclusion, our study demonstrates that although Fgf gain-of-signalling and Hh loss-of-signalling produce similar morphological duplications of the zebrafish ear, they do so via distinct dynamical patterns of gene expression, providing valuable insights into normal anterior otic development. In addition, we determine that *hmx3a*, a gene expressed as an early transcriptional response to both Fgf and Hh manipulation, has a conserved role in correct separation of the sensory maculae within the otic vesicle, and is required—but not sufficient—for normal anterior otic development. We have also shown that our proposed genetic network for zebrafish otic anterior development can be recapitulated with a mathematical model that assumes interactions between a graded extrinsic source of Fgf, a uniform inhibitory influence of Hh, and equipotential competence to adopt an anterior identity at the otic poles. Interactions between these inputs and their downstream targets within the otic tissue (*hmx3a, hmx2, pax5* and *fgf* genes) lead to correct anteroposterior patterning in the developing zebrafish ear. The model will be a useful framework for further elucidation and functional validation of the proposed gene regulatory network required for the acquisition of anterior otic identity in the zebrafish.

## Materials and Methods

### Animals

Adult zebrafish (*Danio rerio*) were kept in circulating water at 28.5°C with a 14-hour light/10-hour dark cycle. The wild-type line used was AB; mutant alleles were *hmx3a^SU3^* (this work; see below for details), and *smo ^hi1640Tg^* (Chen et al., 2001); transgenic lines were *Tg(dusp6:d2EGFP)* (Molina et al., 2007), *Tg(hsp70:fgf3)* (Lecaudey et al., 2008), *Tg(hsp70:fgf8a)*^×*17*^ (Millimaki et al., 2010) and *Tg(hsp70:hmx3a)* (this work; see below for details). The *Tg(hsp70:fgf3)* line was maintained on a *mitfa^w2/w2^* background to reduce pigmentation. Embryos were staged as described (Kimmel et al., 1995) and incubated at 28.5°C in E3 (5 mM NaCl, 0.17 mM KCl, 0.33 mM CaCl_2_, 0.33 mM MgSO_4_, 0.0001% methylene blue), unless otherwise indicated.

### Heat shock

Embryos were cultured in E3 at 28.5°C prior to heat shock. For heat shock, embryos from either a cross between two hemizygous transgenic carriers, or an outcross between a transgenic carrier and a wild-type, were transferred to 25 ml of preheated E3 in a Falcon tube and incubated at 39°C for 30 minutes, unless otherwise indicated. Embryos were then returned to their original plates of E3, which had been preheated to 33°C during the heat shock, and incubated for a further 30 minutes at 33°C. Plates were then returned to 28.5°C and incubated until embryos reached the desired stage for fixation. In heat-shock experiments with mixed batches of transgenic and non-transgenic embryos, a transgenic genotype was confirmed by expression of tdTomato in *Tg(hsp70:hmx3a)* embryos or abnormal shape of the yolk extension in *Tg(hsp70:fgf3)* embryos, in addition to analysis of the phenotypes described in the text.

### Cyclopamine treatment

Embryos were treated in 12-well plates (3 ml total volume; ≤30 embryos per well) at 28.5°C with InSolution Cyclopamine, *V. californicum* (Calbiochem). Chorions were punctured with a sterile hypodermic needle prior to treatment to improve compound penetration. After treatment, embryos were washed twice in E3 before either being fixed or incubated in E3 before fixation later. Vehicle-only controls consisted of a volume of the solvent (ethanol) equivalent to that used in the highest experimental treatment concentration. Embryos from the same batch (siblings) were randomly allocated into control and treatment groups.

### In situ hybridisation

Embryos were dechorionated and fixed in 4% paraformaldehyde overnight at 4°C. In situ hybridisation was carried out as described (Thisse and Thisse, 2008). For most experiments, at least 25 embryos (biological replicates) were stained in any given batch. Where relevant, numbers of embryos with the phenotype of interest and total number in the batch (e.g. 29/30) are shown directly on the figure panels (see figure legends for details). Analysis of gene expression via in situ hybridisation is not quantitative, but we have chosen markers that give a clear and robust qualitative response to changes in signalling pathway activity. We have used information from these spatial expression patterns to infer parameters for the mathematical model (see below). Where appropriate, we have measured the spatial extent of expression along the medial side of the otic vesicle in a dorsal view using ImageJ.

### Generation of a template for the *fgf3* 3’ UTR-specific in situ hybridisation probe

The 3’ UTR of *fgf3* was amplified from wild-type (AB strain) genomic DNA in a nested PCR, incorporating the T7 promoter, using the following primers: F1 TCTCTTGACACAGATGGAGATCC, R1 AATATACAAAGTACTCCTGATTGCA; F2 AAGGCCACTGAGAGTCCAAAA, T7-R2 TAATACGACTCACTATAGGGCAGTAGCCTATCACATGTACGT. Each PCR was run for 30 cycles with an annealing temperature of 53°C.

### Generation of the *hmx3a^SU3^* mutant allele

The single guide RNA (sgRNA) targeting *hmx3a* was designed using CHOPCHOP (Labun et al., 2016; Montague et al., 2014). The sgRNA DNA template was generated using the cloning-free method of Gagnon and colleagues (Gagnon et al., 2014). The template was transcribed and purified using the standard protocols of the MEGAshortscript T7 kit (AM1354, Thermo Fisher Scientific). sgRNA was resuspended in 40 µl of sterile water and the concentration and purity measured using spectrophotometry, before aliquoting for storage at −80°C. To make *Cas9* mRNA, *pCS2-nls-zCas9-nls* plasmid DNA (Jao et al., 2013) was digested with *NotI* and purified by phenol:chloroform extraction, before being transcribed and purified using standard protocols of the mMESSAGE mMACHINE SP6 kit (AM1340, Thermo Fisher Scientific). The resultant mRNA was resuspended, assayed and stored as for the sgRNA. The single cell of one-cell stage AB wild-type embryos was injected with 2 nl of a mixture of 200 ng/µl sgRNA + 600 ng/µl *nls-ZCas9-nls* mRNA. Founders were identified by high resolution melt analysis, using the following primers: PMA F: CGAATGCTAATTTGGCCTCTATTACT and PMA R: TTTTGTTGTCGTCTTCATCGTCC, and Precision Melt Supermix for High Resolution Melt (HRM) Analysis (172-5112, Bio-Rad), performed on a CFX96 Touch System (1855195, Bio-Rad), equipped with Precision Melt Analysis Software (1845025, Bio-Rad). Amplification data were generated using the following program: 95.0°C for 3 minutes, followed by 45 cycles of 95.0°C for 15 seconds, 60.0°C for 20 seconds and 70.0°C for 20 seconds. Melt data were generated using the following program: 65.0°C for 30 seconds, 65.0°C–95.0°C at an incremental rate change of 0.2°C, held for 5 seconds each step, 95.0°C for 15 seconds. Stable F1 heterozygous fish were confirmed by sequencing. All subsequent genotyping was performed by PCR, using the primers F: TGGCAAAGTGACACGACCAG and R: GAGAACACCGTGCGAGTTTTC, Taq DNA Polymerase (M0320S, NEB) and the PCR program: (94.0°C for 2 minutes, 35 cycles of: 94.0°C for 30 seconds, 64.9°C for 30 seconds and 72.0°C for 30 seconds, followed by a final extension at 72.0°C for 2 minutes). The *hmx3a^SU3^* allele is a 69 bp insertion, flanked on either side by 2-base mismatches. The insertion introduces a premature stop codon at nucleotides 352-354 of the edited coding sequence. The insertion in the mutant allele can be distinguished by performing gel electrophoresis on a 2% TBE agarose gel (100V for 40 minutes). The wild-type allele generates a 331 bp product, compared to the 400 bp mutant allele product.

### Generation of the *Tg(hsp70:hmx3a)* line

The zebrafish *hmx3a* cDNA sequence (RefSeq NM_131634.2), including the complete open reading frame, endogenous Kozak sequence and 15 bp of 3’ UTR, was cloned into a *Tol2*-containing *ubi:tdtomato* destination vector, flanked by a 5’ *hsp70* promoter and a 3’ SV40 late polyadenylation signal sequence, using the Tol2kit (Kwan et al., 2007) (Invitrogen). 50 ng of this construct were injected into one-cell stage embryos together with 50 ng of *in vitro*-transcribed transposase RNA. Injected embryos (G0) were raised to adulthood, and their progeny (F1) screened for expression of the tdTomato marker. F1 embryos with positive expression were raised to adulthood to generate a stable *Tg(hsp70:hmx3a)* transgenic line. Progeny were tested by in situ hybridisation after heat shock to check for misexpression of the *hmx3a* transgene.

### Phalloidin staining

Embryos were fixed in 4% PFA overnight, washed in PBS (3×10 minutes) and permeabilised in 2% Triton-X100 (Sigma) for 3-4 days at 4°C. Following further washes in PBS (3×5 minutes), embryos were stained with FITC-phalloidin (1:20; Sigma) or Alexa Fluor 647-phalloidin (1:100; Thermo) in PBS (overnight, 4°C). Embryos were washed in PBS (3×60 minutes), dissected in PBS and mounted in Vectashield (Vectorlabs) prior to confocal imaging.

### Microscopy, photography and image processing

Live and fixed embryos were imaged on either an Olympus BX51 or a Zeiss Axio Imager M1 compound microscope using brightfield, DIC and epifluorescence optics as appropriate, and CellB or Axiovision image acquisition software, respectively. For confocal imaging, either a Nikon A1 or a Zeiss LSM 710 confocal microscope were used. For fluorescent imaging requiring large fields of view, a Zeiss Axio Zoom.V16 stereomicroscope with Zen acquisition software was used. The movie and associated still images shown in the supplementary material for Fig. 7 were acquired with a Zeiss Z.1 light-sheet microscope. Sample drift was corrected using the Manual Drift Correction plugin within FIJI (Fiji Is Just ImageJ) (Schindelin et al., 2012). FIJI was used for all image processing. Figure panels were assembled using Adobe Photoshop 2015.5.0. All dorsal views (except in the supplementary material for Fig. 7) are shown with anterior to the top; lateral views show anterior to the left.

### Statistical analysis

Statistical analyses were performed using GraphPad Prism version 7.0c for Mac OSX (GraphPad software, La Jolla California USA, www.graphpad.com). See figure legends for details.

### Mathematical model

Full information for generation of the mathematical model, including a list of parameters used, is given in Figure 8—Supplemental File 1.

## Acknowledgements

We thank the Gilmour lab for the *Tg(hsp70:fgf3)* line, the Riley lab for the *Tg(hsp70:fgf8a)* line, and Michael Tsang and aquarium staff at University College London for the *Tg(dusp6:d2EGFP)* line. We are grateful to Whitfield lab members for discussion, Sarah Burbridge, Montserrat Garcia Romero, Ginny Grieb and Leslie Vogt for excellent technical support, and the Sheffield and Lewis lab Aquarium Teams for expert fish care.

## Author contributions

RDH: Conceptualisation, investigation, formal analysis, methodology, validation, visualisation, preparation of figures, contribution to writing the manuscript

SJE, KEL: Generation and analysis of the *hmx3a^SU3^* mutant allele, preparation of figures, contribution to writing the manuscript

SB, MM, NvH: investigation, visualisation

NAMM: Supervision, design and generation of the mathematical model, preparation of figures, contribution to writing the manuscript

TTW: Conceptualisation, funding acquisition, project administration, supervision, investigation, validation, formal analysis, visualisation, preparation of figures, writing the manuscript

## Statement on ethical issues

All animal work in the Whitfield lab was covered by licencing from the UK Home Office. All zebrafish experiments conducted in the Lewis lab were approved by the Syracuse University Institutional Animal Care and Use Committee (IACUC).

---

**Figure 1—Supplemental file 1.**
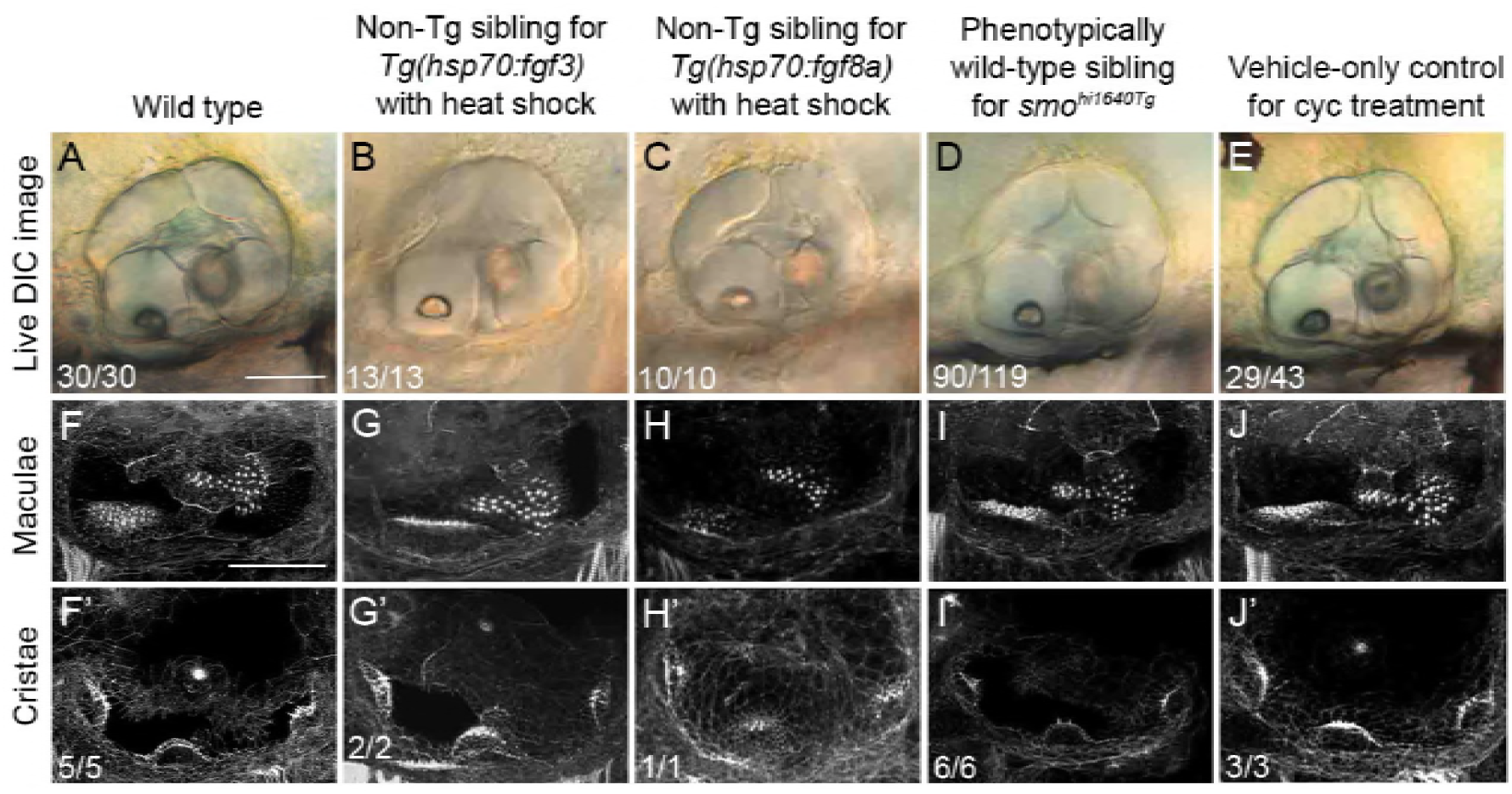
Normal ear development in control embryos for experimental manipulations of Fgf and Hh signalling. **(A–E)** Differential interference contrast (DIC) images of ears in live embryos at 3 dpf (72 hpf). **(F–J’)** Confocal images of FITC-phalloidin stains, revealing stereociliary bundles on sensory hair cells in the maculae (F–J) or cristae (F’–J’), as shown in Fig. 1. The first column (‘Wild type’) repeats column 1 of Fig. 1 for comparison. Subsequent columns show representative images of controls for the experiments shown in Fig. 1. All ears shown were patterned normally, although views and focal planes differ slightly. All ears were of normal size and had two normally-positioned otoliths (A–E), had two maculae of normal size, shape and position in the ear (F–J), and three cristae (F’–J’). Lateral views; anterior to the left. Cyc, cyclopamine. Scale bar in A, 50 μm (applies to A–E); scale bar in F, 50 μm (applies to F–J’).

**Figure 1—Supplemental file 2.**
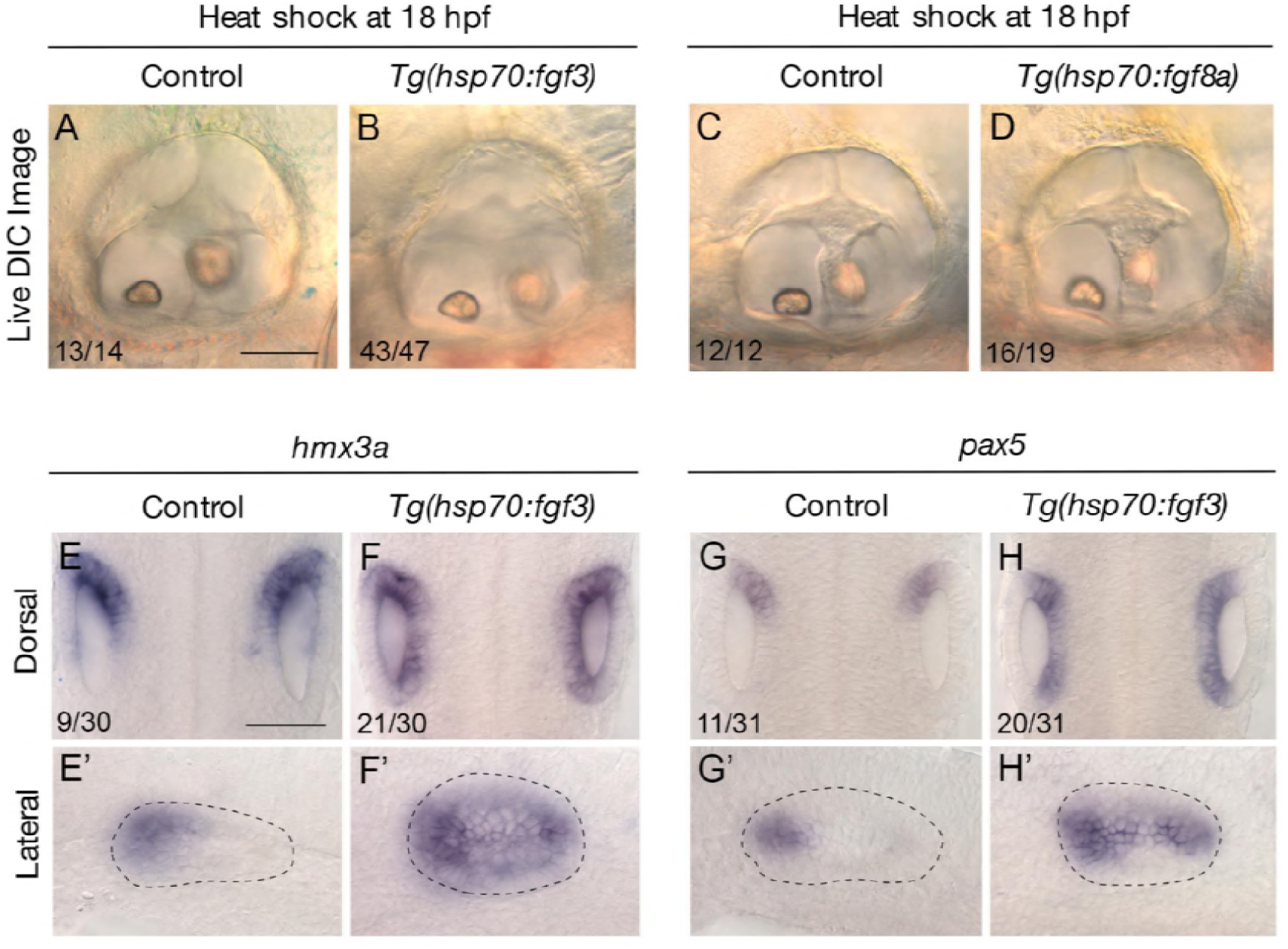
Effects of late mis-expression of *fgf* genes on otic patterning. **(A–D)** Differential interference contrast (DIC) images of ears in live embryos at 3 dpf (72 hpf); lateral views with anterior to the left. Control embryos are non-transgenic siblings subjected to the same heat-shock treatment at 18 hpf. Representative phenotypes are shown; numbers of embryos showing the phenotype are indicated on each panel. Note the relatively normal size and shape of the ears after heat shock in transgenic animals. The focal plane for all panels is at the level of the anterior otolith; note that the posterior otolith (out of focus) is positioned dorsomedially, relative to the anterior otolith, in both control and transgenic ears. **(E–H’)** In situ hybridisation for *hmx3a* (E–F’) and *pax5* (G–H’) at 22.5 hpf. Note the expansion of expression for both markers after heat shock of transgenic animals. E–H are dorsal views showing both ears; E’–H’ are lateral views with anterior to the left. Scale bar in A, 50 μm (applies to A–D); scale bar in E, 50 μm (applies to E–H’).

**Figure 2—Supplemental file 1.**
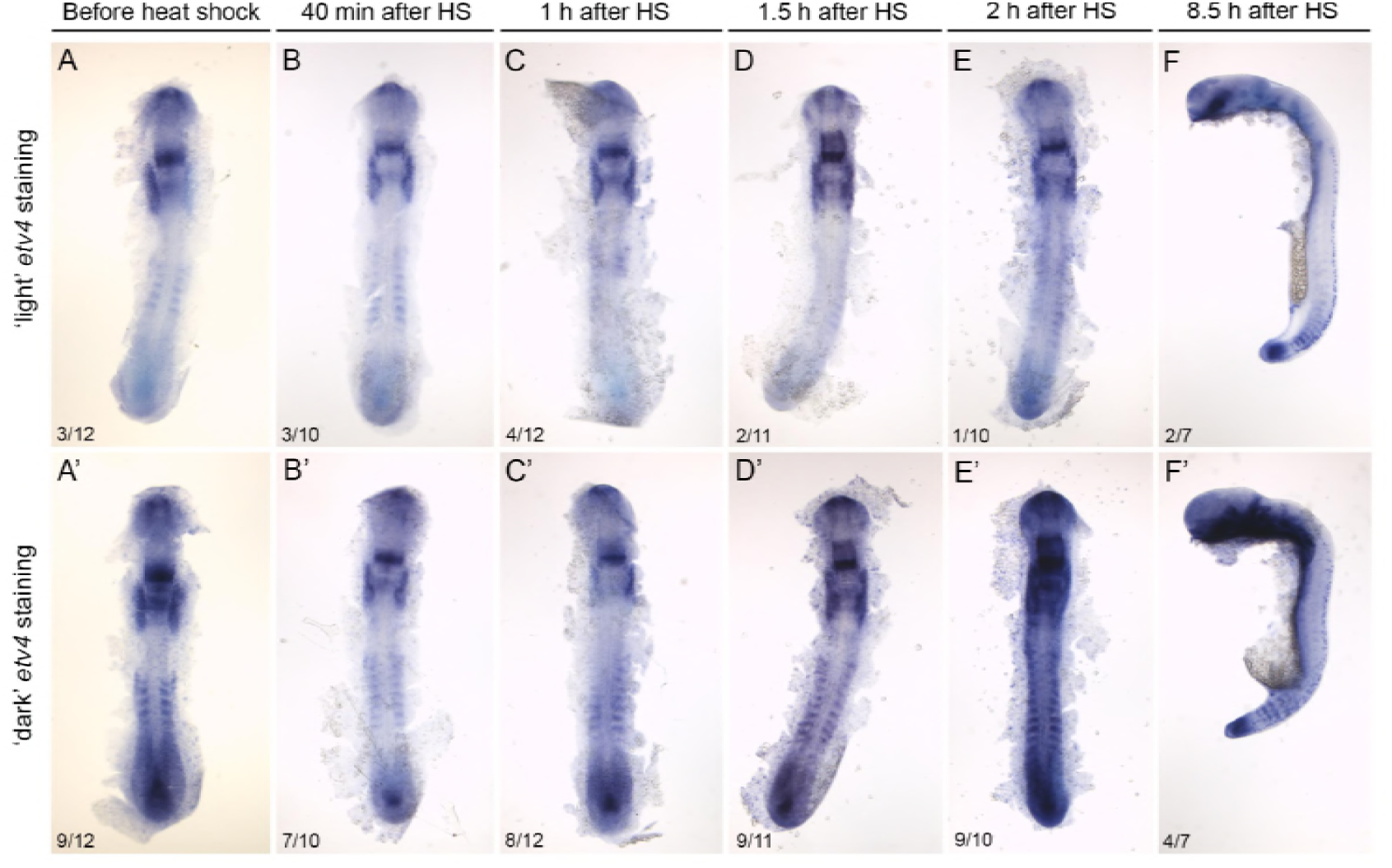
Time-course of expression of *etv4* mRNA after heat shock in *Tg(hsp70:fgf3)* embryos. Embryos were heat-shocked for 30 minutes at 39°C at 14 hpf (the 10-somite stage), fixed at various times after the onset of heat shock as shown (top), and processed for in situ hybridisation for *etv4.* All embryos were stained and photographed using bright field optics under identical conditions, and scored as having ‘light’ or ‘dark’ expression (presumed transgenic and non-transgenic embryos, respectively; 75% of the batch was expected to be transgenic). Number of embryos with the phenotype shown and total number in the batch are shown directly on the panels (e.g. 3/12). Expression levels at the 10-somite stage before heat shock (A, A’) were very variable, possibly due to leaky expression of the transgene, but corresponded to the published spatial pattern of expression (Thisse and Thisse, 2004). Robust, systemic up-regulation of *etv4* was seen in embryos 2 hours after heat shock (E, E’). This persisted 8.5 hours after heat shock; here, 4/5 presumed transgenic embryos with abnormal morphology also had strong *etv4* expression (F’). Morphology was normal in presumed non-transgenic siblings showing the endogenous expression pattern of *etv4* (F). F, F’ show lateral views; all other panels are dorsal views of flat-mounted embryos.

**Figure 2—Supplemental file 2.**
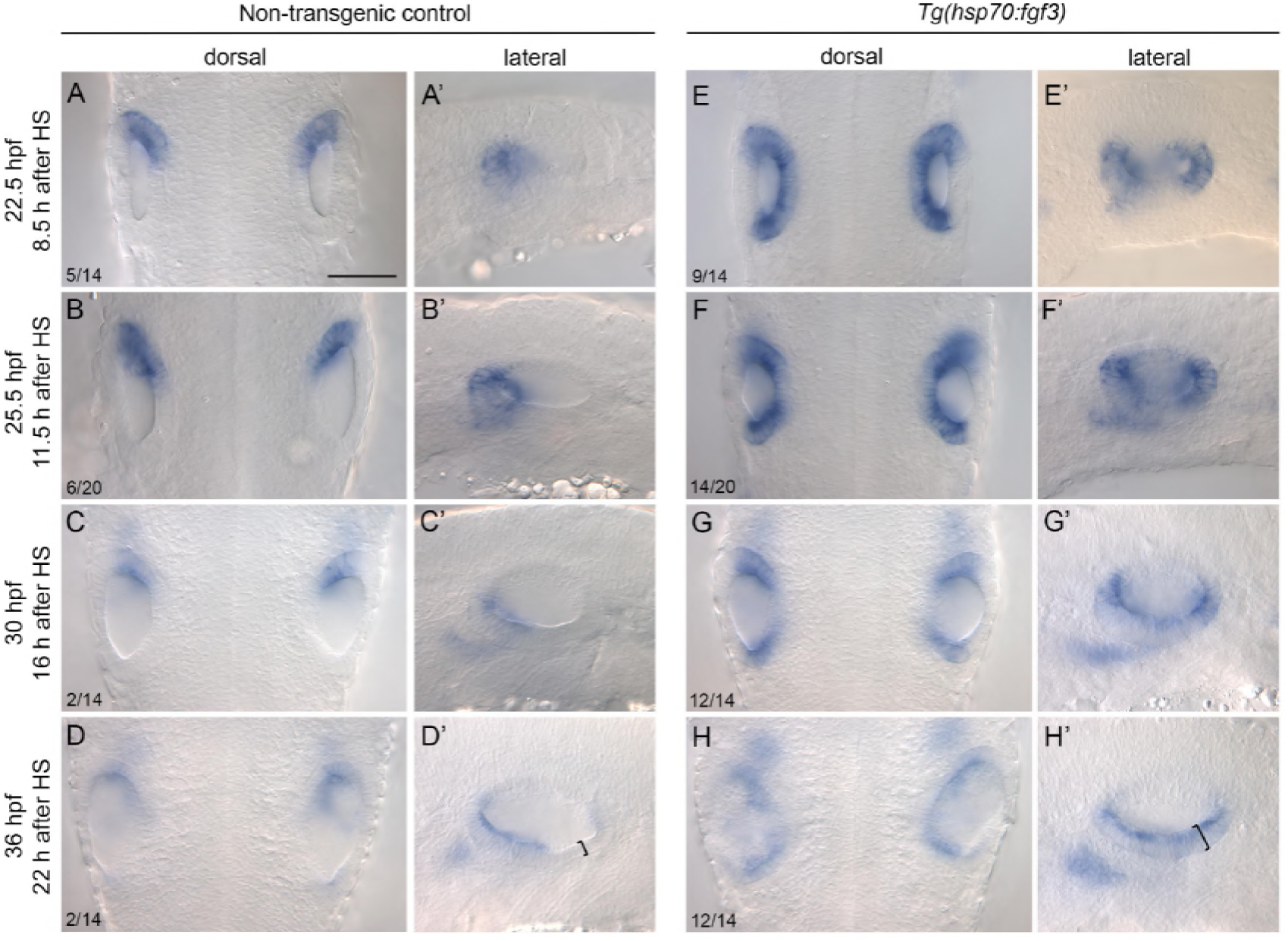
Detailed time-course of expression of *hmx2* after early *fgf3* mis-expression. In situ hybridisation of otic expression of *hmx2* in *Tg(hsp70:fgf3)* embryos following a 30-minute heat shock (HS) at the 10-somite stage (14 hpf). Controls (A-D’) were sibling nontransgenic embryos subjected to the same heat shock. Numbers in the dorsal view panels indicate the number of embryos with the phenotype shown and total number (e.g. 5/14) from a mixed batch of transgenic and non-transgenic embryos in each pair of panels; 75% of the batch was expected to be transgenic. The first and last rows are biological replicates of data shown in Fig. 2. Note the weakening of expression in central medial otic epithelium by 25.5 hpf in transgenic embryos. By 36 hpf, in a lateral view, *hmx2* is expressed throughout the ventral floor of the otic vesicle in transgenic embryos, associated with a thicker epithelium in posterolateral regions (D’, H’, brackets). All dorsal views show anterior to the top; all lateral views show anterior to the left. Scale bar in A, 50 μm (applies to all panels).

**Figure 2—Supplemental file 3.**
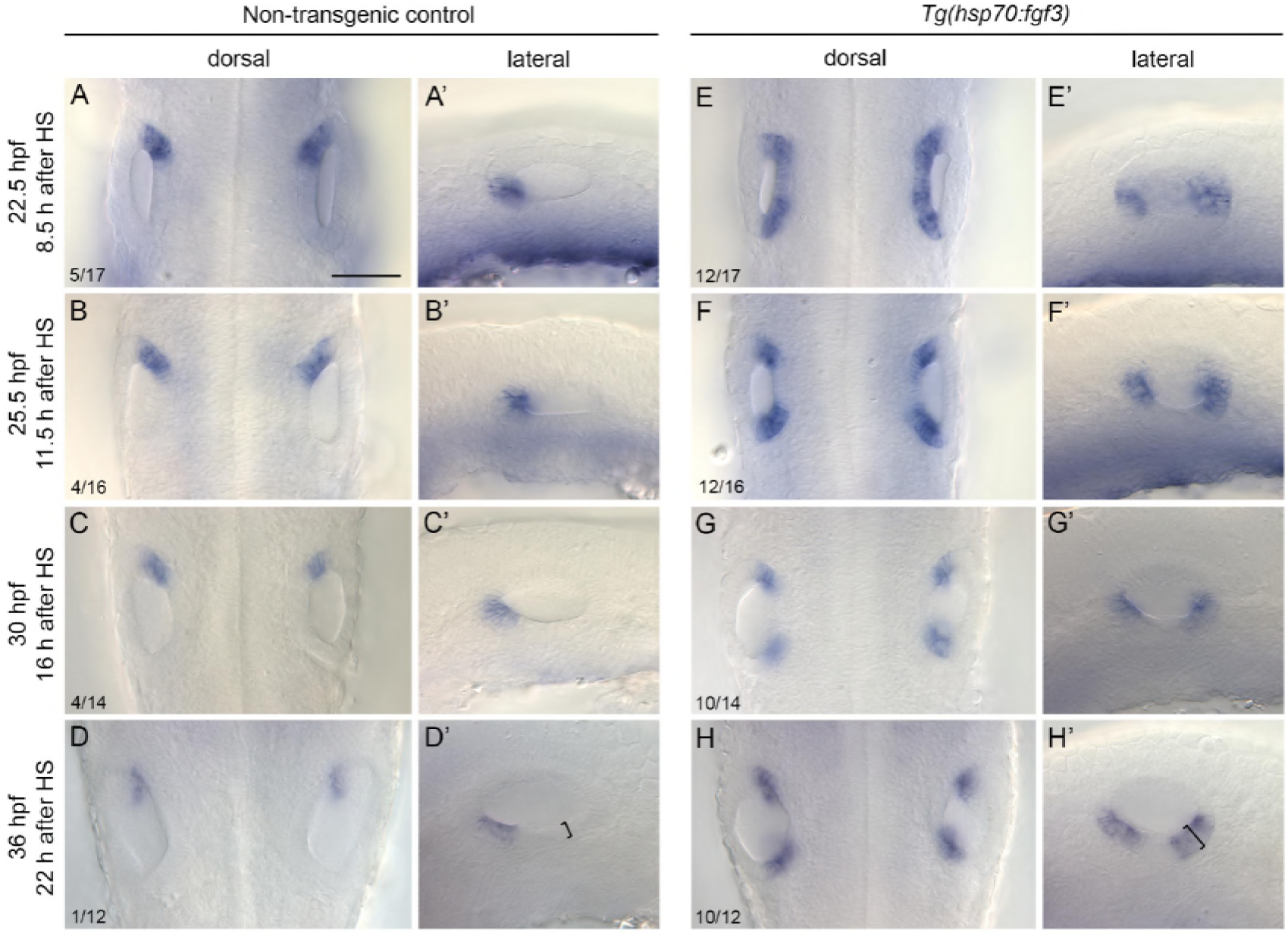
Detailed time-course of expression of *pax5* after early *fgf3* mis-expression. In situ hybridisation of otic expression of *pax5* in *Tg(hsp70:fgf3)* embryos following a 30- minute heat shock (HS) at the 10-somite stage (14 hpf). Controls (A-D’) were sibling nontransgenic embryos subjected to the same heat shock. Numbers in the dorsal view panels indicate the number of embryos with the phenotype shown and total number (e.g. 5/17) from a mixed batch of transgenic and non-transgenic embryos in each pair of panels; 75% of the batch was expected to be transgenic. The first and last rows are biological replicates of data shown in Fig. 2. Note that the ectopic expression in transgenic embryos has already resolved into two domains by 25.5 hpf, and two discrete domains persist at 36 hpf. At 36 hpf, in a lateral view, the ectopic domain of *pax5* is associated with a thicker epithelium (D’, H’, brackets). One embryo at 36 hpf was unable to be scored. All dorsal views show anterior to the top; all lateral views show anterior to the left. Scale bar in A, 50 μm (applies to all panels).

**Figure 3—Supplemental file 1.**
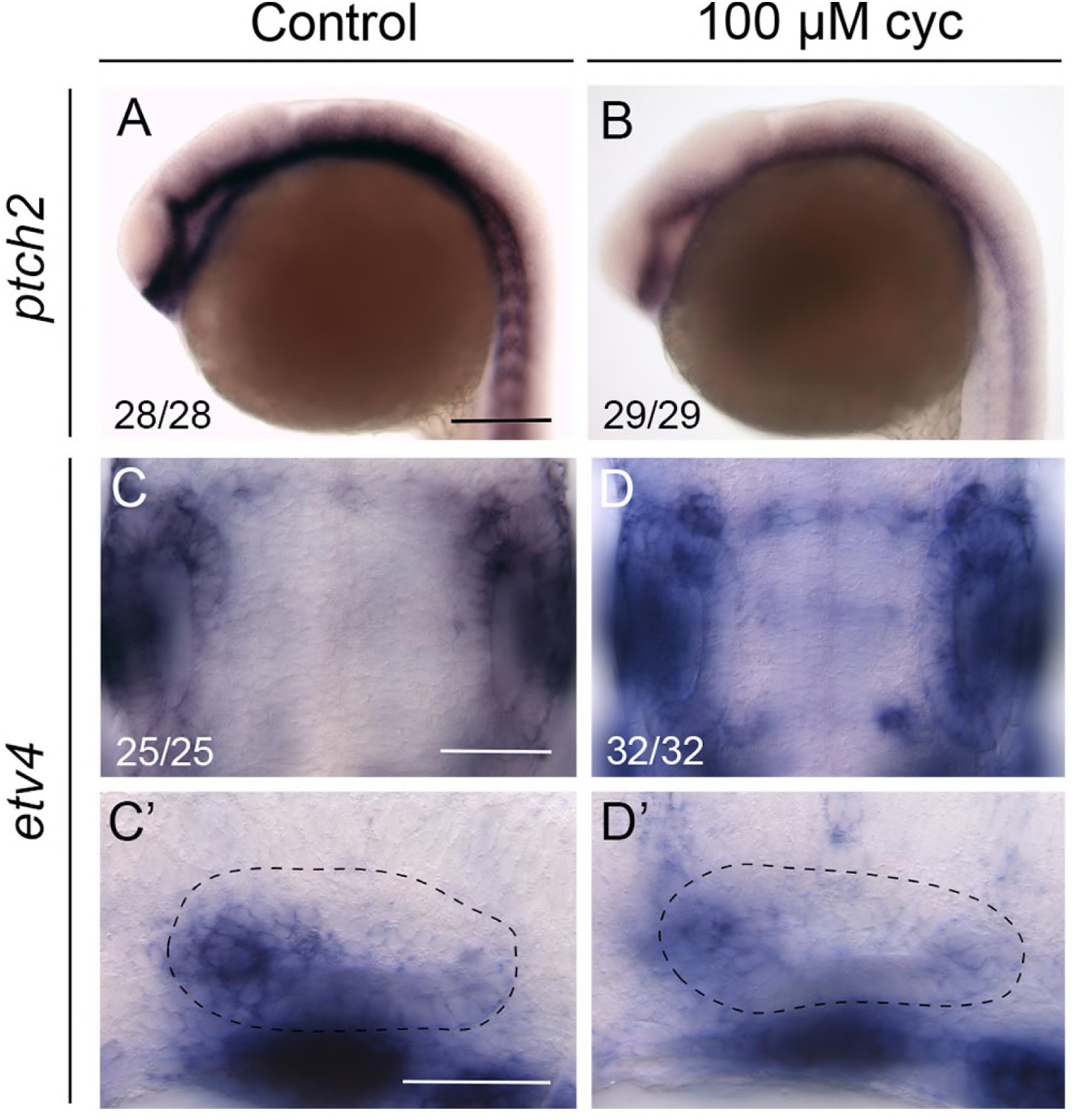
Expression of target genes of the Hh and Fgf pathways after cyclopamine treatment. **(A, B)** In situ hybridisation for the Hh pathway target gene *ptch2* at 22.5 hpf, after treatment from 14.5 hpf with vehicle (ethanol) only (A) or 100 μM cyclopamine (cyc; B). The head and yolk of the embryo are shown. Lateral views; anterior to the left. Expression is reduced but not abolished in cyclopamine-treated embryos (B). **(C–D’)** In situ hybridisation for the Fgf target gene *etv4* at at 22.5 hpf, after treatment from 14.5 hpf with vehicle (ethanol) only (C, C’) or 100 μM cyclopamine (cyc; D, D’). There are no major changes to the otic expression of *etv4* at this stage after cyclopamine treatment. C, D show dorsal views of the two otic vesicles with anterior to the top; C’, D’ are lateral views of the otic vesicle with anterior to the left. Scale bar in A, 200 μm (applies to B); scale bar in C, 50 μm (applies to D); scale bar in C’, 50 μm (applies to D’).

**Figure 3—Supplemental file 2.**
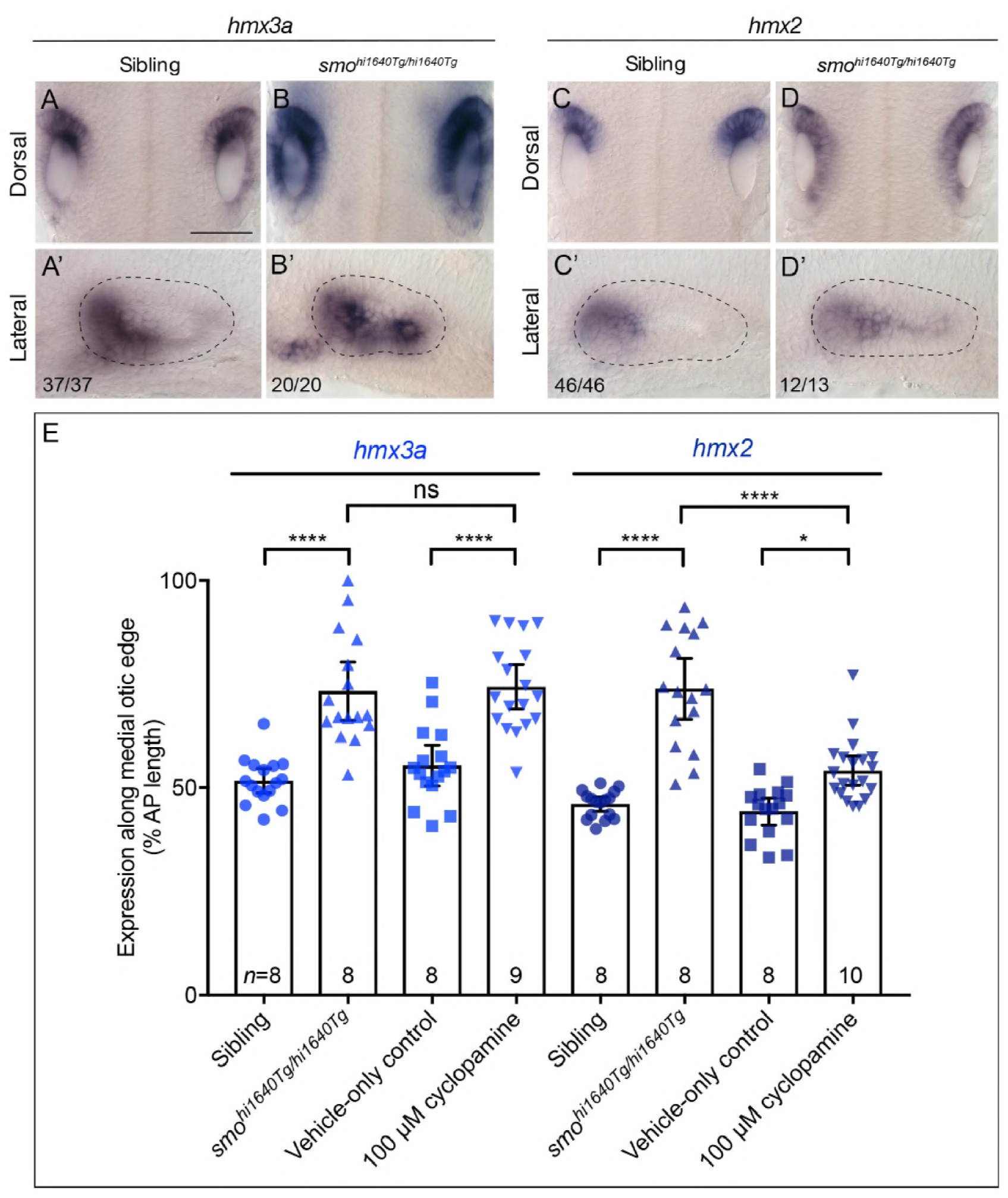
Otic expression of *hmx* genes after genetic or pharmacological inhibition of Hh signalling. **(A–D’)** In situ hybridisation for *hmx* genes at 22.5 hpf in the otic vesicles of phenotypically wild-type and *smo^hl1640T9/hl1640T9^* mutant embryos. In the mutant, otic expression extends posteriorly in a graded fashion at this stage. A–D show dorsal views of the two ears, with anterior to the top; A’–D’ show lateral views of the otic vesicle with anterior to the left. Scale bar in A, 50 μm (applies to A–D); scale bar in A’, 50 μm (applies to A’–D’). **(E)** Measurements of the extent of in situ hybridisation stain along the medial edge of the otic vesicle at 22.5 hpf, expressed as a percentage of the length of the medial edge of the otic epithelium. The expression domain of both *hmx* genes extends posteriorly in *smo^hl1640T9/hl1640T9^* mutant embryos, and after treatment with 100 μM cyclopamine from 14 hpf. One-way ANOVA with Šídák’s post-test correction for multiple comparisons: \*\*\*\**p*<0.0001; \**p*=0.0119; ns, nonsignificant (*p*=0.9998). Error bars represent the 95% confidence interval for the mean. *n* indicates number of embryos; two ears were measured for each embryo, with each ear measurement shown as a separate data point.

**Figure 5—Supplemental file 1.**
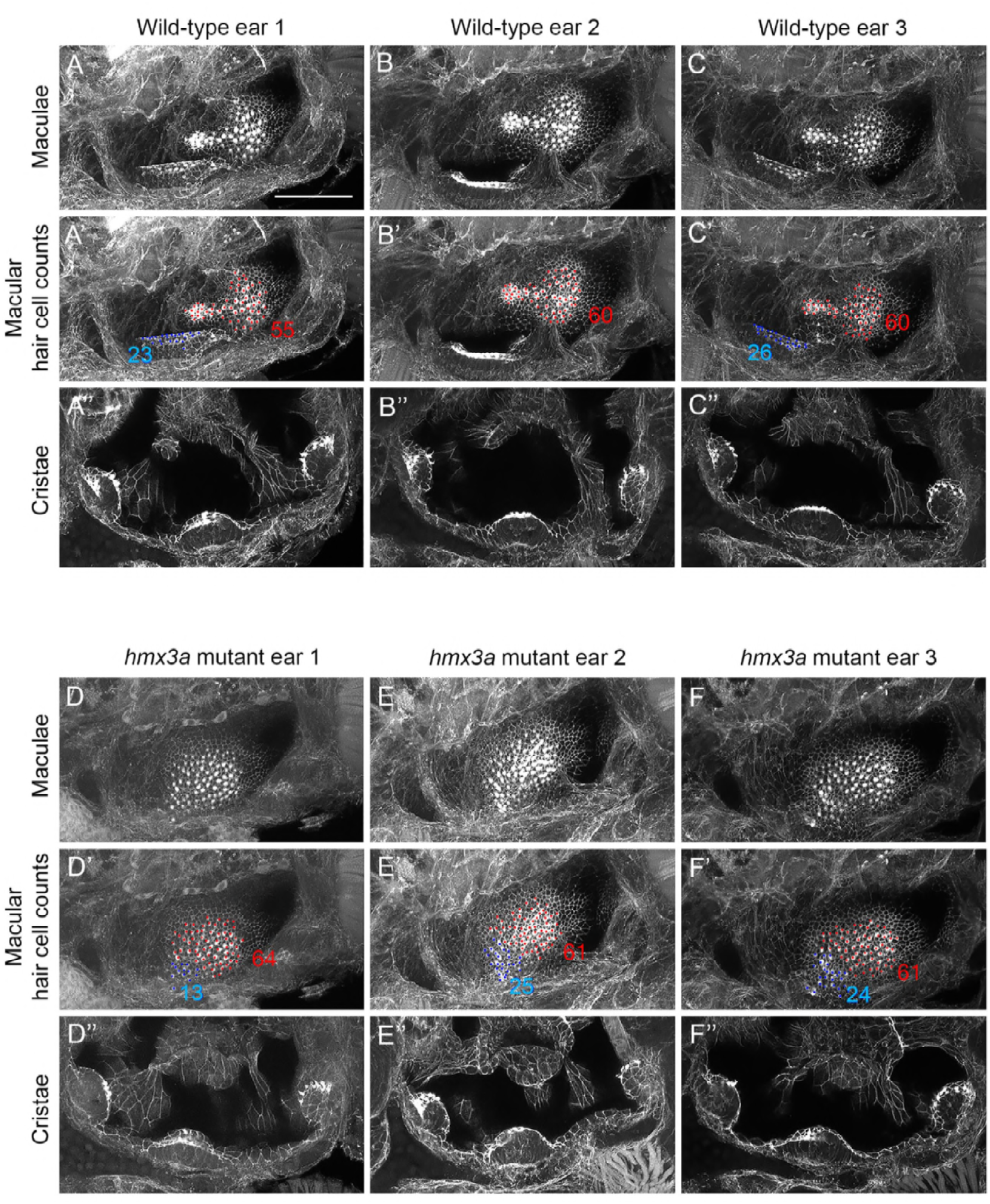
Maculae are fused, but cristae are normal, in *hmx3a^SU3/SU3^* mutant ears. **(A–F’’)** Confocal images of FITC-phalloidin-stained ears at 3 dpf (72 hpf); maximum intensity projections of selected z-stacks. Three ears were imaged for each genotype; note the fused maculae, but normal presence of three cristae, in each of the mutant ears (D–F’’). The middle row of panels in each set is a duplicate of the panels above, with counts for visible hair cells in the anterior macula (blue) and posterior macula (red). The distinction between the anterior and posterior parts of the fused macula in panels D’–F’ was estimated based on hair cell position. Anterior macula counts in the wild-type samples are likely to be underestimates, as only some of the hair cells were visible in this orientation. It was not possible to distinguish any individual hair cells in the anterior macula in wild-type ear 2 (B’). The number of ears imaged was too small to draw firm conclusions about any changes in hair cell number in either macula. Note that the panels for wild-type ear 3 and *hmx3a^SU3/SU3^* mutant ear 3 are reproduced in Fig. 5. Scale bar, 50 μm (applies to all panels).

**Figure 5—Supplemental file 2.**
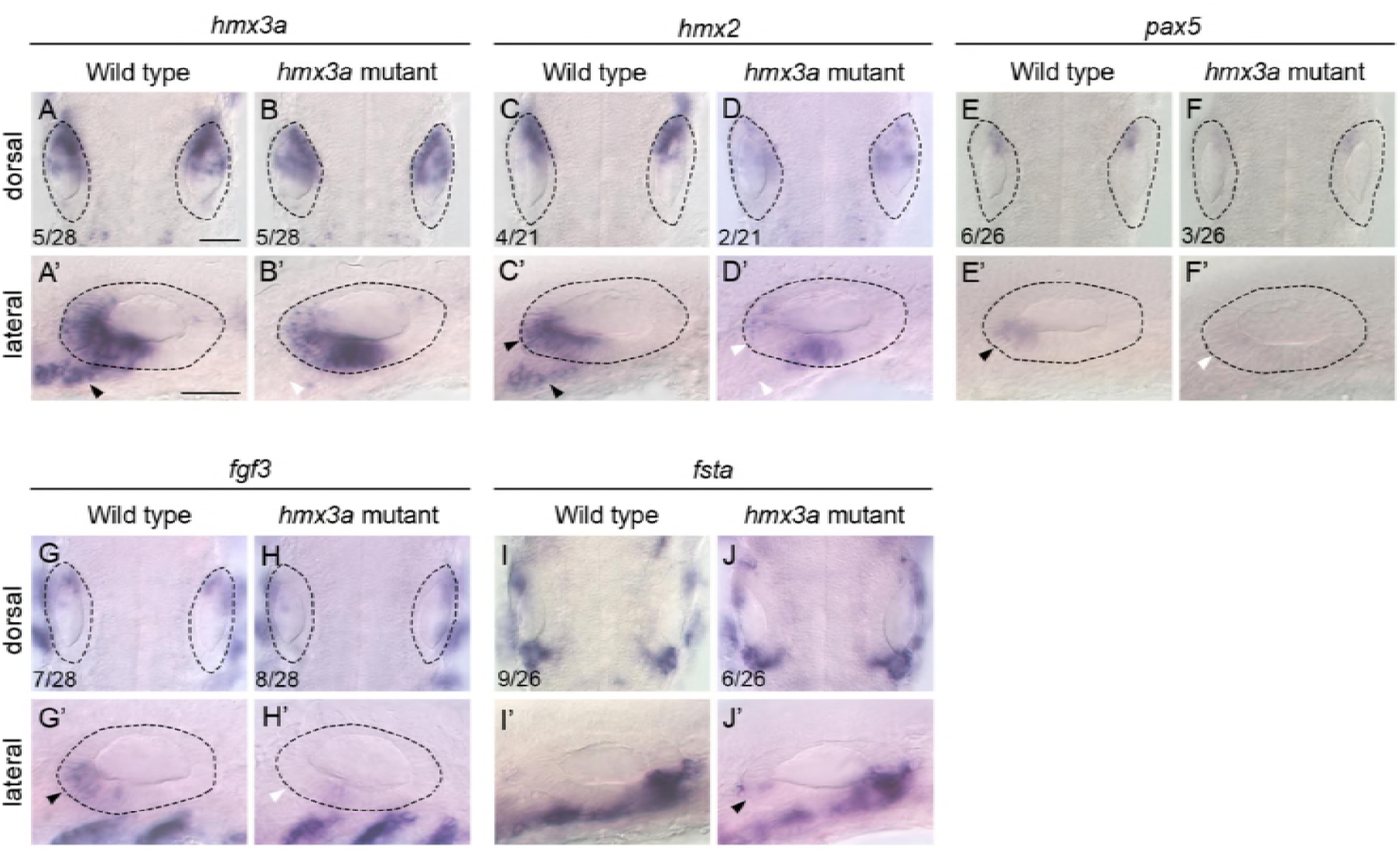
Expression of otic markers at 27–30 hpf in *hmx3a^SU3/SU3^* mutants. (**A–H’**) Reduction in expression of anterior otic markers in the otic vesicle of *hmx3a^SU3/SU3^* mutants at 27 hpf. Black arrowheads mark expression domains in wild-type ears and in presumptive neuroblasts anteroventral to the ear that are reduced or missing in mutants (white arrowheads). (**I–J’**) Expression of the posterior otic marker *follistatin-a (fsta)* in the otic vesicle at 30 hpf. Although weak *fsta* expression was detected in anterior otic epithelium in mutants (J’, black arrowhead), levels were not above the natural variation seen in wild-type siblings. Numbers in panels A–J indicate numbers of embryos genotyped as either wild type or homozygous mutant that showed the representative expression patterns illustrated. Scale bar in A, 50 μm (applies to A–J); scale bar in A’, 50 μm (applies to A’–J’).

**Figure 6—Supplemental file 1.**
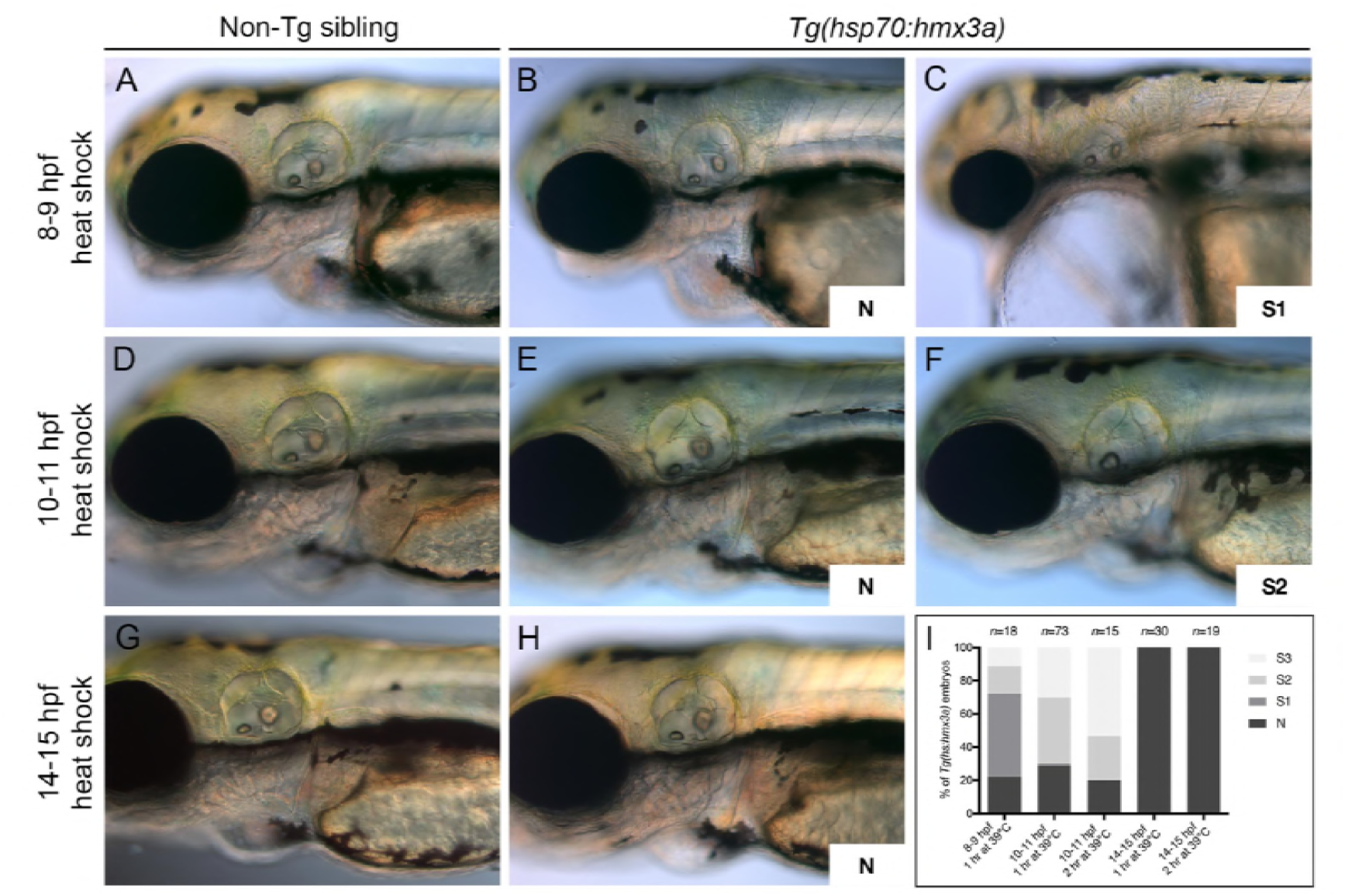
Development of embryos and otic phenotypes after mis-expression of *hmx3a* at different time points. **(A-H)** Transgenic *Tg(hsp70:hmx3a)* and non-transgenic sibling embryos were heat-shocked for 1 hour at the times indicated, and grown on to 3 dpf for examination of general morphology and any otic phenotype. Representative examples are shown. **(I)** Graphical representation of the results from 1- and 2-hour heat shocks. Abbreviations: N, normal positioning of both anterior (ventrolateral) and posterior (posteromedial) otoliths; S1 - anterior otolith in wild-type ventrolateral position but posterior otolith more ventrally positioned; S2 - single otolith in one ear with two wild-type positioned otoliths in the contralateral ear; S3 – single otolith in both ears; *n*, number of embryos.

**Figure 7—Supplemental file 1.**
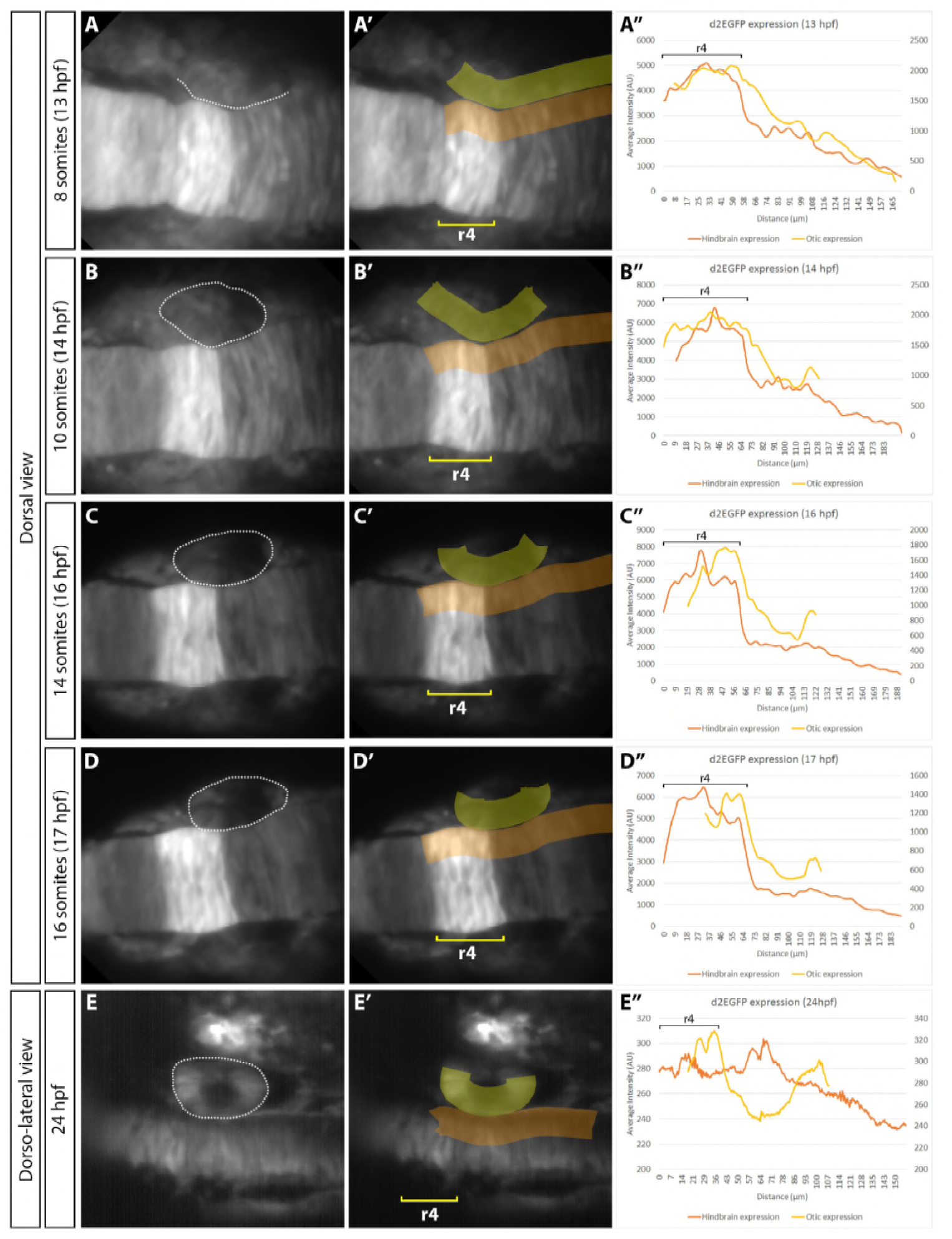
Expression of a fluorescent reporter for Fgf activity in the hindbrain and otic placode. (A–E”) Light-sheet imaging and fluorescence measurements of a representative *Tg(dusp6:d2EGFP)* embryo from the 8-somite stage to the 16-somite stage (A–D), and a second representative embryo at 24 hpf (E). A dorsal view of the hindbrain and right-hand otic region (dotted outline) is shown; anterior is to the left. Rhombomere 4 (r4) expression is bright during the 8–16-somite stages. Expression levels were averaged over a 28.7 μm-wide band, over 20 *z*-sections at intervals of 1 μm, both in the hindbrain posterior to the r3/r4 boundary (orange; left-hand scale on graph) and otic region (yellow; right-hand scale on graph). Position of the otic region was estimated by tracing the dataset backwards from a stage when the otic vesicle was evident. At 24 hpf, expression of d2EGFP in r4 decreased, whereas the otic vesicle had high Fgf activity at both the anterior and posterior poles (E–E’’). The measurements at 24 hpf were taken from a different embryo using a lower laser power, resulting in different arbitrary units on the graph (E’’). At earlier stages, expression of d2EGFP in the otic region was much lower than that in the rhombomeres, but a graded expression (higher at the anterior, lower at the posterior) was evident between the 8- and 16- somite stages (A–D’’). The images in A–D’ were taken from the dataset shown in Supplemental Movie 1.

**Supplemental movie 1**

Time-lapse movie of a representative *Tg(dusp6:d2EGFP)* embryo from the 8-somite stage to the 16-somite stage, used to generate panels A–D’ in Figure 7—Supplemental file 1. Images were captured by light-sheet microscopy every 5 minutes for 4 hours (from the 8- to the 16-somite stage). Individual time points were manually drift corrected. An average intensity projection of the fluorescent signal is shown (produced from 20 *z*-slices encompassing the otic region). A dorsal view is shown; anterior is to the top left. Scale bar, 50 µm.

**Figure 8—Supplemental file 1. Details of the mathematical model.** See appendix.

## Details of the mathematical model

### Representation of otic tissue

For the purposes of modelling gene expression in the developing otic tissue between 14 and 36 hours post fertilisation (hpf), we represent the medial side of the otic tissue as a one-dimensional array of cells. Distance along this array—represented by the variable *x*—is measured in percentage length along the anterior-posterior (AP) axis. At the stages studied, the length of the medial side is approximately 100 µm.

### Competence to express *fgf*

Our data suggest that competence to express *fgf* in the otic tissue (*fgf_i_*) begins at around 14 hpf (see Fig. 7B) and is strongest at the poles. We therefore assume that this competence can be represented by a function of the form

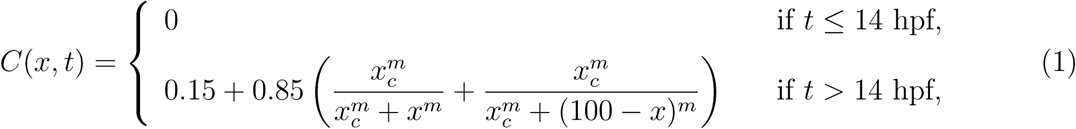

where 0 ≤ *x* ≤ 100 is percentage length along the AP axis, *x_c_* is a measure of the extent of the polar competence regions, and *m* is a measure of the sharpness of the boundaries between regions of high and low competence. In our model simulations, we assume *x_c_* = 20% AP length and *m* = 5, giving the competence profile shown in Fig. S1A.

### Extrinsic Fgf expression

We assume that rhombomere 4 of the hindbrain acts as the main source of extrinsic Fgf signalling; both *fgf3* and *fgf8a* are expressed here at the time of initial otic anteroposterior patterning (Maves et al., 2002). Further support is provided by analysis of mutants for *mafba* (Kwak et al., 2002) and *hnf1ba* (Lecaudey et al., 2007). These two genes code for transcription factors expressed in the hindbrain and are required for restriction of *fgf3* expression to rhombomere 4. In the *mafba^−/−^* and *hnf1ba^−/−^* mutants, posterior expansions of *fgf3* expression in the hindbrain correlate with expansions or duplications of otic anterior markers similar to those we see in *Tg(hsp70:fgf3)* embryos after heat shock.

We assume that the extrinsic Fgf3 protein spreads away from the cells in which it is produced, resulting in a graded expression profile in the neighbouring otic tissue. In support of this, we show in Fig. 7 – Supplemental File 1 that a reporter of Fgf signalling activity is expressed in a decreasing gradient in the otic region, with highest activity in the tissue neighbouring rhombomere 4. Up to time *t* = 18 hpf, we assume that the distribution of rhombomeric Fgf protein in the otic tissue is given by

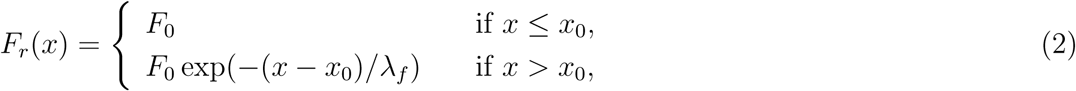

where *F_r_* (*x*) is the Fgf protein concentration (in arbitrary units) in the otic tissue at AP position *x*, 0 ≤ *x* ≤ *x*_0_ is the region of overlap between rhombomere 4 and the otic tissue, *F*_0_ is the maximum protein concentration, and λ*_f_* is the effective “diffusion wavelength” of Fgf. The resulting concentration profile is shown in Fig. S1B for *F*_0_ = 18.5, *x*_0_ = 30% AP axis, and λ*_f_* = 15% AP axis (approximately 15 µm at the stages studied). *fgf* expression in rhombomere 4 decreases significantly at around 18 hpf (Maves *et al.*, 2002). Up until this time, the Fgf concentration profile in the otic tissue is given by Eq. (2). At times later than 18 hpf, we assume linear degradation of the Fgf protein. The Fgf concentration at time *t* is therefore given by

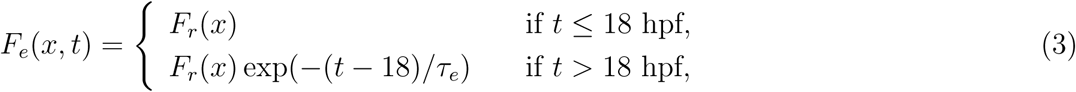

where *τ_e_* is the half-life (in hours) of the Fgf protein.

**Figure S1:**
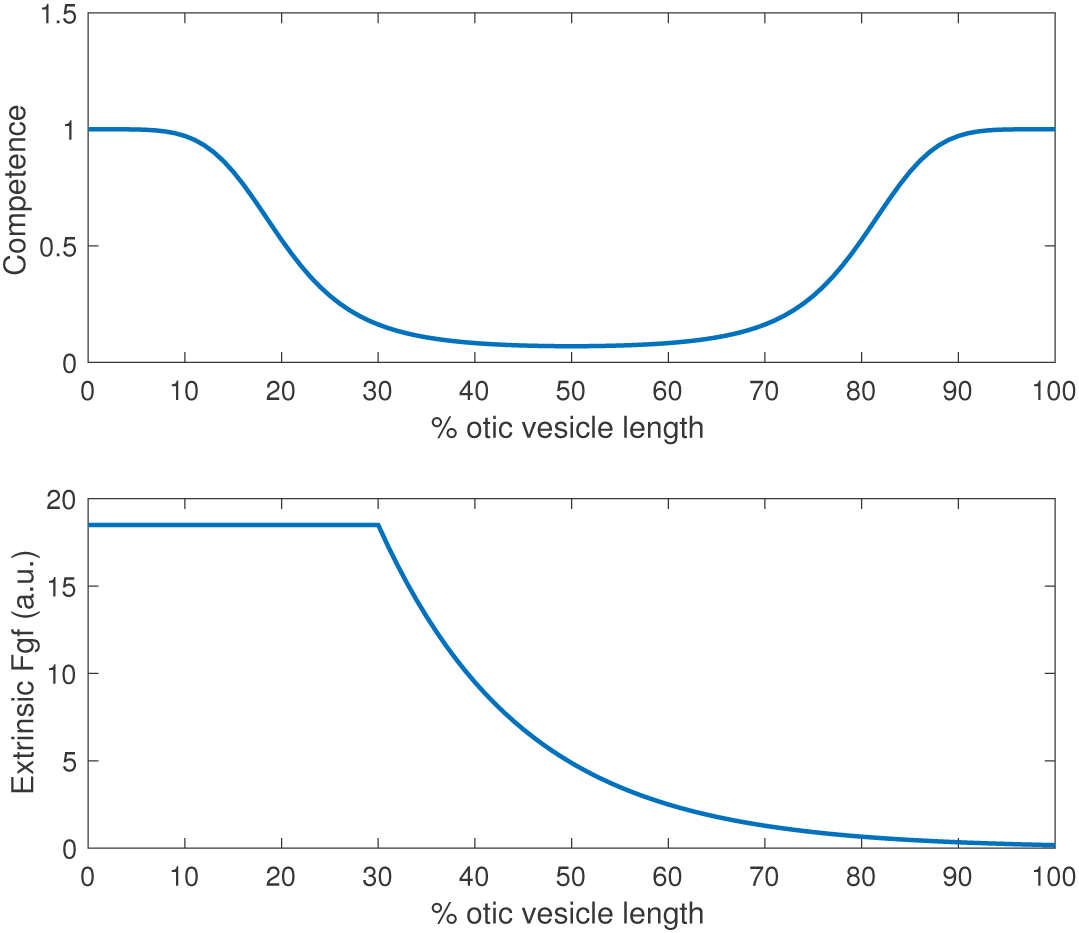
(A) Assumed distribution of competence for endogenous *fgf* expression in the otic tissue with *x_c_* = 20% AP length and *m* = 5 (see Eq. (1)); (B) Assumed distribution (up until 18 hpf) in the otic tissue of Fgf protein produced in rhombomere 4 (see Eq. (2)). *F*_0_ = 18.5, *x*_0_ = 30% AP axis, λ*_f_* = 15% AP axis.

### Regulated gene expression in the otic tissue

Fgf protein originating from rhombomere 4 initiates a temporal sequence of spatially patterned gene expression in the otic tissue. Based on the inferred interactions summarised in Fig. 7A, we represent transcription and translation using a system of coupled differential equations as follows:

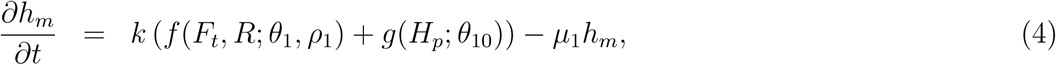

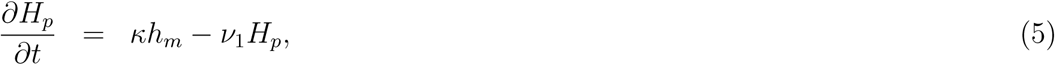

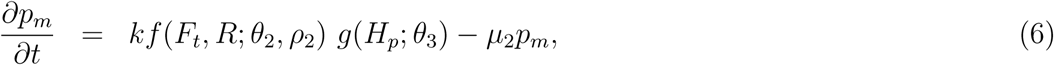

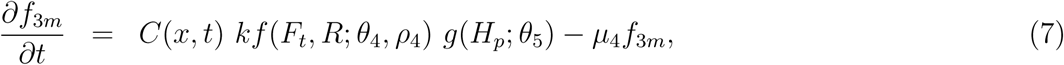

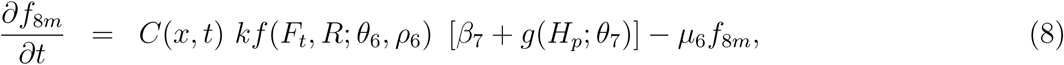

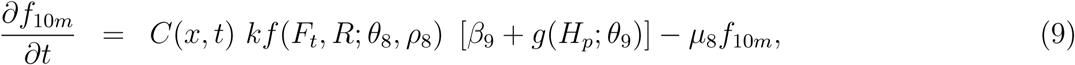

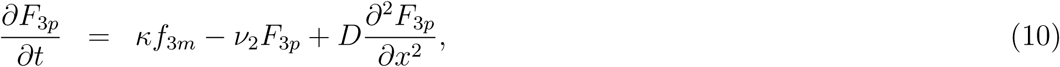

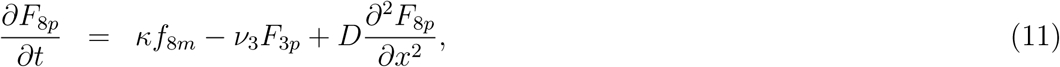

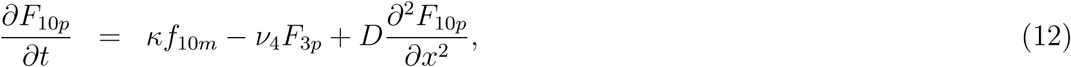

where

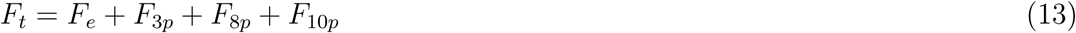

represents the total amount of Fgf protein in the otic tissue and the transcription regulation functions are given by increasing sigmoid (Hill) functions of the general form:

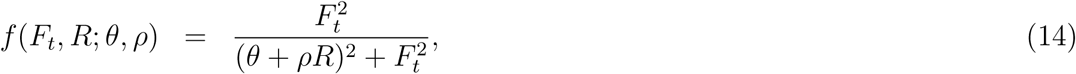

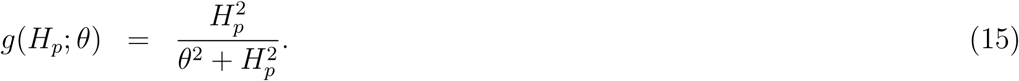

In these functions, the parameter θ represents the activation threshold — the concentration of activator required to achieve a half-maximal rate of transcription. The effect of Hh attenuation is to increase the threshold in Eq. (14) by an amount *ρR*, where *R* is a measure of the amount of Hh signalling in the otic tissue, and *ρ* is the relative attenuation strength for each gene. Hh attenuation thus reduces the rate of transcription resulting from a given concentration of Fgf.

The meaning of all model variables and parameters is summarised in Tables S1–S3.

In the model, expression of all genes is activated by the total amount of Fgf protein (both extrinsic and that produced within the otic tissue) and attenuated by Hh protein. Expression of *hmx3a*, *pax5*, *fgf3*, *fgf8a* and *fgf10a* is additionally activated by Hmx3a protein.

**Table S1:** Definition of model variables

| *Variable* | *Meaning* |
| --- | --- |
| $h_m$ | *hmx3a* mRNA concentration |
| $H_p$ | Hmx3a protein concentration |
| $p_m$ | *pax5* mRNA concentration |
| $f_{3m}$ | *fgf3* mRNA concentration |
| $f_{8m}$ | *fgf8a* mRNA concentration |
| $f_{10m}$ | *fgf10a* mRNA concentration |
| $F_{3p}$ | Fgf3 protein concentration |
| $F_{8p}$ | Fgf8a protein concentration |
| $F_{10p}$ | Fgf10a protein concentration |
| $F_e$ | Extrinsic Fgf protein concentration |
| $F_t$ | Total Fgf protein concentration |
| $R$ | Hh protein concentration |
| $C(x, t)$ | Competence to express *fgf* |
| $F_e(x, t)$ | Extrinsic Fgf protein |

Simulations of the model were performed with the parameter values listed in Tables S2 and S3 on a discrete spatial domain comprising 100 spatial cells (with the diffusion terms in Eqs. (10)–(12) represented by a simple finite difference scheme), with zero-flux boundary conditions at the anterior and posterior poles of the otic vesicle. Simulations covered the time period 10–36 hpf, with initial values of all intrinsic mRNA and protein variables set to zero. The resulting spatiotemporal mRNA expression patterns are shown in Fig. 8, for three conditions: wild type (1st column), heat shock induction of *fgf3* (2nd column), and inhibition of Hh signalling by cyclopamine treatment (3rd column). The simulation protocols for the latter two conditions are described below. Fig. S2 shows the spatial profiles of mRNA and total Fgf protein expression for the three conditions at 22.5 hpf and 36 hpf.

Transcription and translation rates for all endogenous genes and proteins have been set to 1, so all expression levels are expressed in arbitrary units. Half-lives of mRNA and protein have been set to reflect the observed dynamics of expression patterns. For example, the *pax5* mRNA half life is set to be low (0.5 hrs) to reflect the fact that *pax5* mRNA expression induced in the middle part of the otic vesicle by heat shock Fgf3 protein is lost by 36 hpf. The *hmx3a* mRNA half life is also set to be low (0.5 hrs) to reflect the early onset of *hmx3a* expression. In contrast, the *fgf* mRNA half lives are set to be higher to reflect the later onset of their expression. The Fgf protein diffusion coefficient is set to be low in order to avoid Fgf protein produced in the anterior and posterior poles “flooding” the otic vesicle. The short half lives of the Fgf proteins also contribute to the restriction of Fgf proteins to the poles. Indeed, Fgf protein diffusion can be omitted from the model without affecting the dynamics of the mRNA expression patterns.

The transcription regulation parameters, which reflect the level of expression at which regulating proteins effect regulation of their targets, were chosen with reference to the expression levels achieved by each protein in the model (shown in Fig. S3). For example, the threshold for regulation of *hmx3a* expression by Fgf protein (*θ*_1_) is the primary determinant of the extent of the anterior expression domain of *hmx3a* mRNA.

**Table S2:** Production, degradation and diffusion parameters

| *Parameter* | *Value* | *Description* |
| --- | --- | --- |
| $k$ | 1 a.u. per hr | Maximum transcription rate |
| $\kappa$ | 1 a.u. per mRNA per hr | Translation rate |
| $k_{HS}$ | 100 a.u. per hr | *fgf3* heat shock transcription rate |
| $\mu_1$ | $\ln 2/0.5 \text{ hr}^{-1}$ | *hmx3a* degradation rate |
| $\mu_2$ | $\ln 2/0.5 \text{ hr}^{-1}$ | *pax5* degradation rate |
| $\mu_4$ | $\ln 2/3 \text{ hr}^{-1}$ | *fgf3* degradation rate |
| $\mu_6$ | $\ln 2/6 \text{ hr}^{-1}$ | *fgf8a* degradation rate |
| $\mu_8$ | $\ln 2/4 \text{ hr}^{-1}$ | *fgf10a* degradation rate |
| $\nu_1$ | $\ln 2/1.5 \text{ hr}^{-1}$ | Hmx3a degradation rate |
| $\nu_2$ | $\ln 2/0.5 \text{ hr}^{-1}$ | Fgf3 degradation rate |
| $\nu_3$ | $\ln 2/0.5 \text{ hr}^{-1}$ | Fgf8a degradation rate |
| $\nu_4$ | $\ln 2/0.5 \text{ hr}^{-1}$ | Fgf10a degradation rate |
| $D$ | $50 \mu\text{m}^2 \text{ hr}^{-1}$ | Fgf diffusion coefficient |
| $F_0$ | 18.5 | Extrinsic Fgf amount (a.u.) |
| $R$ | 15 | Hh attenuation strength |

### Simulation of heat shock induction of *fgf3a* expression and cyclopamine treatment

To simulate heat shock induction of *fgf3* expression, we include additional variables to represent *fgf3* mRNA and Fgf3 protein produced from the *fgf3* transgene. We assume that transcription from the transgene starts at 14 hpf and terminates at 14.5 hpf. Because of the time taken for the resulting mRNA and protein to decay, the effects of the heat shock on target gene expression extend beyond 14.5 hpf.

To simulate treatment of embryos with cyclopamine (a chemical inhibitor of Hh signalling) at 14 hpf, we assume that a consequent reduction in the Hh-dependent antagonism of Fgf-dependent transcription (the variable *R* in the model equations) does not begin until 15 hpf. In this way, we represent the time taken for a reduction in Hh signalling to feed through to a reduction in the intracellular effectors of Hh signalling. We further assume that the inhibitory term *R* decays exponentially for *t* > 15 hpf, with a half-life of 3 hrs.

**Figure S2:**
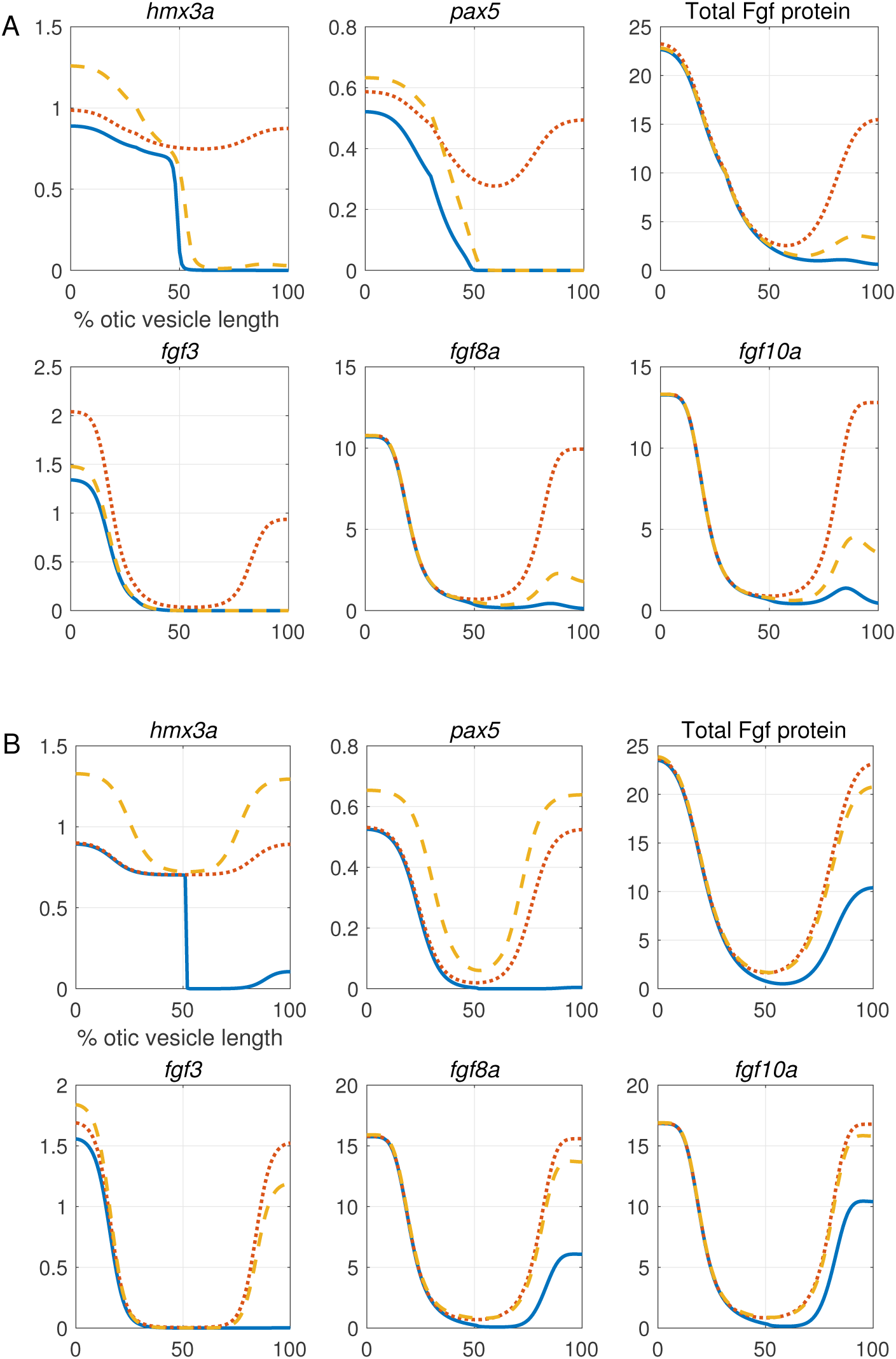
Spatial expression profiles of model variables at 22.5 hpf (A) and 36 hpf (B). In each panel, the solid blue curve represents the wild type, the dotted red line represents heat shock induction of *fgf3*, and the dashed orange line represents cyclopamine treatment (inhibition of Hh signalling). mRNA profiles (*hmx3a*, *pax5*, *fgf3*, *fgf8a*, *fgf10a*) are measured as transcripts per cell; Fgf protein is measured as protein molecules per cell, and “Total Fgf protein” represents *F_t_* — the sum of all Fgf proteins (intrinsic and extrinsic).

**Figure S3:**
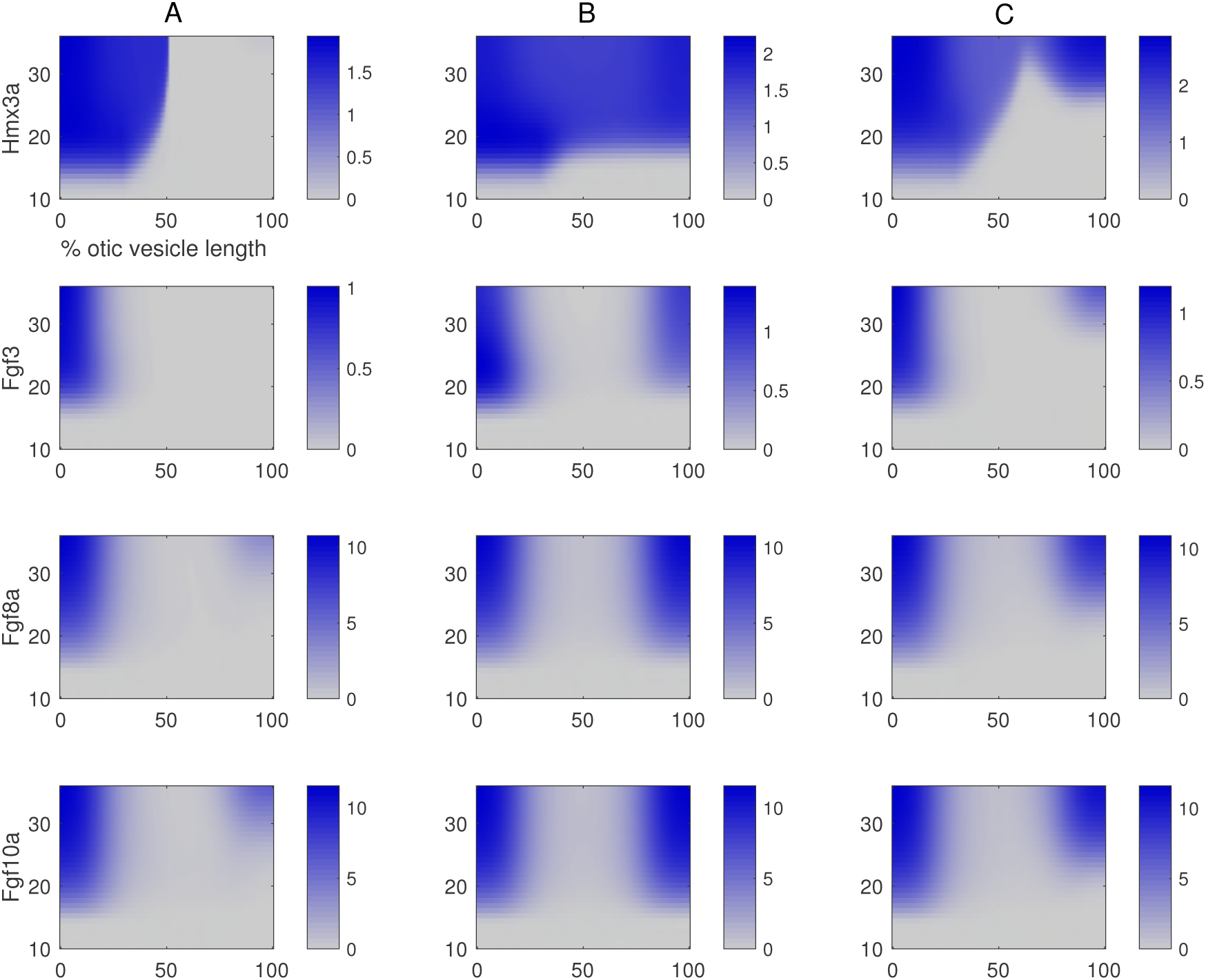
Spatio-temporal expression profiles of Hmx3a, Fgf3, Fgf8a and Fgf10a protein model variables in simulations of wild type (column A), heat shock induction of *fgf3* (column B), and cyclopamine treatment (column C)

**Table S3:** Transcription regulation parameters

| *Parameter* | *Value* | *Description* |
| --- | --- | --- |
| $\theta_1$ | 10 | Fgf-*hmx3a* threshold |
| $\theta_2$ | 5 | Fgf-*pax5* threshold |
| $\theta_3$ | 0.68 | Hmx3a-*pax5* threshold |
| $\theta_4$ | 25 | Fgf-*fgf3* threshold |
| $\theta_5$ | 0.83 | Hmx3a-*fgf3* threshold |
| $\theta_6$ | 1 | Fgf-*fgf8a* threshold |
| $\theta_7$ | 0.06 | Hmx3a-*fgf8a* threshold |
| $\theta_8$ | 1 | Fgf-*fgf10a* threshold |
| $\theta_9$ | 0.04 | Hmx3a-*fgf10a* threshold |
| $\theta_{10}$ | 0.25 | Hmx3a-*hmx3a* threshold |
| $\rho_1$ | 2 | Hh-*hmx3a* attenuation coefficient |
| $\rho_2$ | 0.4 | Hh-*pax5* attenuation coefficient |
| $\rho_4$ | 0.1 | Hh-*fgf3* attenuation coefficient |
| $\rho_6$ | 0.1 | Hh-*fgf8a* attenuation coefficient |
| $\rho_8$ | 0.05 | Hh-*fgf10a* attenuation coefficient |
| $\beta_7$ | 1 | Hmx3a-independent *fgf8a* production coefficient |
| $\beta_9$ | 2 | Hmx3a-independent *fgf10a* production coefficient |

